# Blood progenitor redox homeostasis through GABA control of TCA cycle in *Drosophila* hematopoiesis

**DOI:** 10.1101/2021.02.23.432543

**Authors:** Manisha Goyal, Ajay Tomar, Sukanya Madhwal, Tina Mukherjee

## Abstract

The importance of reactive oxygen species (ROS) in myeloid cell development and function is well-established. However, a comprehensive understanding of metabolic states controlling ROS levels during hematopoiesis remains elusive. Myeloid-like blood progenitor cells of the *Drosophila* larvae reside in a specialized hematopoietic organ called the lymph gland. We find that these progenitors in homeostasis, utilize TCA to generate ROS. Excessive activation of TCA however raises ROS levels causing them to precociously differentiate and leads to retardation of lymph gland size. Thus, to maintain ROS homeostasis, progenitor cells utilize systemically derived GABA. GABA internalization and catabolism via inhibiting hydroxy prolyl hydroxylase (Hph) activity, promotes pyruvate dehydrogenase kinase enzyme activity (PDK). PDK controls inhibitory phosphorylation of pyruvate dehydrogenase (PDH), the rate-limiting enzyme, connecting pyruvate to TCA cycle and OXPHOS. Thus, by regulating PDK, GABA regulates progenitor TCA activity and ROS levels. In addition to this, GABA-catabolism/Hph axis via Hifα/Sima drives a glycolytic state in progenitor cells. The dual control established by GABA on PDK and Sima maintains progenitor cell metabolism and sustains ROS homeostasis necessary for their development. Taken together, our study demonstrates the metabolic underpinnings of GABA in myeloid ROS regulation and their development, the relevance of which may be broadly conserved.

## Introduction

The utilization of reactive oxygen species (ROS) as a developmental signal in immune-progenitor development and fate decisions is apparent both in vertebrates and invertebrates (Bigarella et al., 2014; Harris et al., 2013; Prieto-Bermejo et al., 2018; Takubo et al., 2013; Tothova et al., 2007; Vincent & Crozatier, 2010). The developmental roles for ROS are reliant on its threshold levels as any aberrant generation of ROS can alter immune stem or progenitor-cell development at the level of their maintenance, differentiation, or function (Dragojlovic-Munther & Martinez-Agosto, 2012; Owusu-Ansah & Banerjee, 2009). Thus, mechanisms underlying ROS homeostasis during hematopoietic development comprise an integral component of redox signalling. In this context, an understanding of metabolic programs that enable blood-progenitor cells to co-ordinate their ROS levels are still limited and forms the focus of our investigation.

*Drosophila* larval blood progenitors akin to the mammalian common-myeloid blood progenitors (CMP), reside in a hematopoietic organ termed the lymph gland. Here, they are maintained in the undifferentiated state via signals emanating from the local niche (also called posterior signalling center, PSC) (Krzemien et al., 2007; Morin-Poulard et al., 2016), differentiating hemocytes and systemic cues derived from the brain and fat body (Banerjee et al., 2019). The signalling cues include Hh (Mandal et al., 2007), wingless (Sinenko et al., 2009), JAK/STAT (Makki et al., 2010), Dpp (Dey et al., 2016), TGF-β (Makhijani et al., 2017) and insulin (Benmimoun et al., 2012). These are conserved developmental pathways also well-established for their control of mammalian hematopoiesis (Clements & Traver, 2013). Additionally, recent studies have highlighted important metabolic contributions in blood progenitor development. Progenitor sensing of metabolites derived either locally, like ROS (Owusu-Ansah & Banerjee, 2009), amino acids (Shim et al., 2012), lipids (Tiwari et al., 2020) and adenosine (Mondal et al., 2011) or systemically from neurons (Madhwal et al., 2020; Shim et al., 2013), have been implicated in governing diverse aspects of progenitor development and function. The lymph gland hematopoietic system therefore with its intrinsic feature of metabolic and signaling requirements, offers a perfect developmental model to gain a comprehensive view of metabolic programs controlling progenitor ROS homeostasis in blood development.

A key source of ROS in cells is carbon cycling or the citric acid cycle (TCA) (Quinlan et al., 2012; Sabharwal & Schumacker, 2014). TCA generates multiple intermediates and also controls mitochondrial oxidative phosphorylation (OXPHOS) leading to ROS generation (Kaplon et al., 2013; Mailloux et al., 2016; Quinlan et al., 2012; Starkov et al., 2004). TCA is driven by the entry of pyruvate, which is an end-product of glycolysis, and is also derived from additional sources in the cellular cytoplasm. Ultimately, it is destined for transport into mitochondria as a master fuel input driving citric acid cycle and oxidative phosphorylation (Gray et al., 2014; Wang et al., 2016). Pyruvate is converted to acetyl CoA via pyruvate dehydrogenase enzyme (PDH), the key enzyme linking glycolysis to TCA. Its activity is regulated by post-translational modification, which involves phosphorylation driven by pyruvate dehydrogenase kinase (PDK) which inactivates PDH. The dephosphorylation event by pyruvate dehydrogenase phosphatase (PDP) activates the enzyme fueling the TCA cycle (Bowker-Kinley et al., 1998; Harris et al., 2002).

Lymph gland blood progenitor cells have been shown to maintain elevated levels of ROS. However, when in excess it causes oxidative stress leading to precocious differentiation and loss of progenitor homeostasis (Dragojlovic-Munther & Martinez-Agosto, 2012; Owusu-Ansah & Banerjee, 2009). Progenitor cells also rely on GABA, which is neuronally derived upon olfactory stimulation and is sensed by blood cells both as a signaling entity and as a metabolite. GABA signaling via GABABR in lymph gland progenitor cells controls their maintenance (Shim et al., 2013) while its utilization as a metabolite regulates progenitor differentiation potential (Madhwal et al., 2020). Intracellularly, GABA catabolism regulates Hifα/Sima protein stability in progenitor cells necessary to maintain progenitor responses to infections by parasitic wasps (Madhwal et al., 2020). The current study explores the importance of GABA metabolic pathway in governing TCA activity in the control of progenitor ROS homeostasis. We find that in homeostatic conditions, the fueling of pyruvate into the TCA-cycle and activation of mitochondrial complex II protein, succinate dehydrogenase (SDH), a key TCA enzyme leads to ROS generation in progenitor cells. However, elevation in TCA activity limits overall growth of the hematopoietic lymph gland tissue and leads to loss of progenitor maintenance. Thus, to maintain TCA activity, progenitor cells adopt the GABA catabolic pathway and limit pyruvate’s entry into the TCA. Specifically, we find that GABA via its end product succinate inhibits hydroxy prolyl hydroxylase (Hph) activity and maintains active PDK function. This suppresses PDH enzymatic activity leading to lower TCA rate and homeostatic control of ROS generation. Furthermore, GABA via Sima stabilization also regulates key glycolytic enzyme, Lactate dehydrogenase (Ldh), known to promote a glycolytic state. Thus, by employing GABA to inhibit Hph function, progenitor cells limit TCA and support glycolytic state which allows these cells to maintain ROS balance and supports their normal development.

## Results

### GABA metabolism in blood progenitor cells controls overall size of the lymph gland

*Drosophila* lymph gland blood progenitor cells internalize GABA via GABA transporter (Gat) and catabolize it into succinate (Fig. 1A, Madhwal et al., 2020). Succinic-semialdehyde dehydrogenase (Ssadh), the last and the rate-limiting step of GABA catabolic pathway (Shelp et al., 1999) is responsible for generation of succinate in progenitor cells. Succinate via inhibition of hydroxy prolyl hydroxylase (Hph) activity, stabilizes Sima protein in progenitor cells. Sima functions orthologous to mammalian Hifα, where prolyl hydroxylase activity of Hph marks Sima protein for proteasomal mediated degradation. In the presence of GABA, blood progenitor cells stabilize Sima protein, whose downstream target lactate dehydrogenase (Ldh) is known to capacitates blood cells with differentiation potential necessary to respond to parasitic wasp-infections (Madhwal et al., 2020).

**Figure 1.**
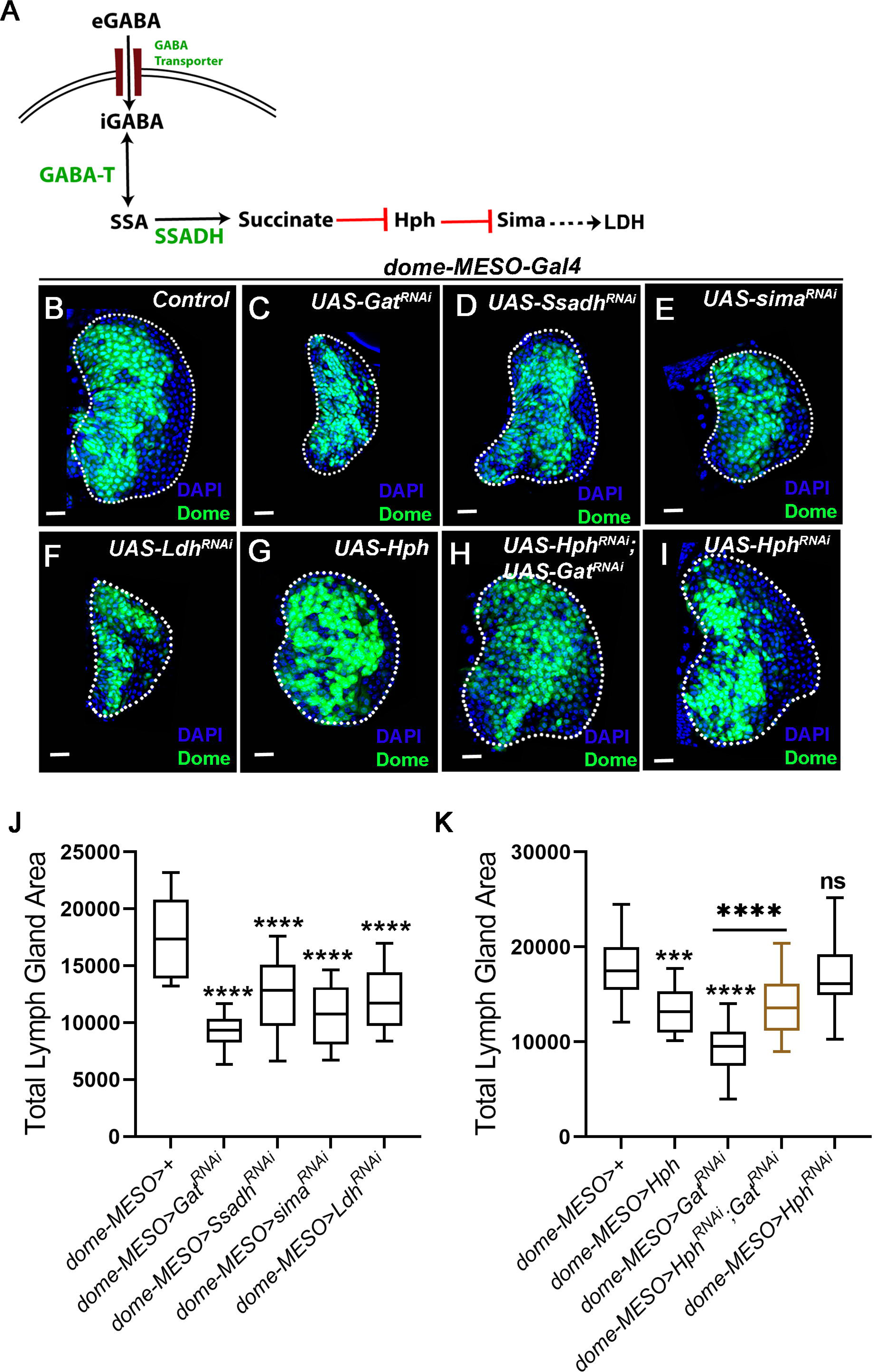
GABA catabolism in *Drosophila* blood progenitor cells controls lymph gland growth. DNA is marked with DAPI (blue). Asterisks mark statistically significant differences (*p<0.05; **p<0.01; ***p<0.001, ****p<0.0001, Mann-Whitney test for **J** and **K** is employed). Lymph glands are demarcated with white dotted line. Scale bar: 20µm. ‘n’ represents number of lymph gland lobes analysed. **(A)** Schematic representation of the GABA-catabolic pathway and downstream Ldh regulation in blood progenitor cells. Uptake of extra cellular eGABA (GABA) via GABA Transporter (GAT) in blood progenitor cells and its intracellular catabolism into succinate by GABA-transaminase (GABA-T) and succinic semi-aldehyde dehydrogenase (SSADH, rate limiting step). Succinate stabilizes Sima protein by inhibiting the activity of Hph enzyme and Sima transcriptionally control *Ldh* expression. **(B-G)** Representative images showing lymph gland growth status, compared to **(B)** control lymph gland (*dome-MESO>GFP/+*), **(C)** expressing *Gat^RNAi^* in progenitor cells (*dome-MESO-Gal4, UAS-GFP; UAS-Gat^RNAi^*) and **(D)** expressing *Ssadh^RNAi^* in progenitor cells (*dome-MESO-Gal4, UAS-GFP; UAS-Ssadh^RNAi^*) leads to reduction in lymph gland size. Similarly, expressing **(E)** *sima^RNAi^* (*dome-MESO-Gal4, UAS-GFP; UAS-sima^RNAi^*), **(F)** *Ldh^RNAi^* (*dome-MESO-Gal4, UAS-GFP; UAS-Ldh^RNAi^*). For quantifications, refer to **J. (G-I)** Over-expressing *Hph* (*dome-MESO-Gal4, UAS-GFP; UAS-Hph*) show reduction in lymph gland size as compared to **(A)** control, expressing *Hph^RNAi^* in *Gat^RNAi^* (*dome-MESO-Gal4, UAS-GFP; UAS-Hph^RNAi^*; *UAS-Gat^RNAi^*) leads to rescue of lymph gland growth defect as compared to **(C)** *Gat^RNAi^*, **(I)** expressing *Hph^RNAi^* in (*dome-MESO-Gal4, UAS-GFP; UAS-Hph^RNAi^*) leads to no change in lymph gland growth as compared to control **(B).** For quantifications, refer to **K. (J)** Quantification for total lymph gland area in *dome-MESO-Gal4, UAS-GFP*/+ (control, n=20), *dome-MESO-Gal4, UAS-GFP*; *UAS-Gat^RNAi^* (n=20, p<0.0001), and *dome-MESO-Gal4, UAS-GFP*; *UAS-Ssadh^RNAi^* (n=20, p<0.0001), *dome-MESO-Gal4, UAS-GFP*; *UAS-sima^RNAi^* (n=20, p<0.0001) and *dome-MESO-Gal4, UAS-GFP*; *UAS-Ldh^RNAi^* (n=20, p<0.0001). (K) Quantifications for total lymph gland area in *dome-MESO-Gal4, UAS-GFP*/+ (control, n=20), *dome-MESO-Gal4, UAS-GFP; UAS-Hph* (n=20, p=0.0002), *dome-MESO-Gal4, UAS-GFP*; *UAS-Gat^RNAi^* (n=20, p<0.0001), and *dome-MESO-Gal4, UAS-GFP; UAS-Hph^RNAi^*; *UAS-Gat^RNAi^* (n=20, p<0.0001, as compared to *Gat^RNAi^*) and *dome-MESO-Gal4, UAS-GFP*; *UAS-Hph^RNAi^* (n=20, p=0.5784).

We observed that loss of components of this pathway (Fig. 1A) from blood progenitor cells led to a significant reduction in the overall size of the lymph gland (Fig. 1B-F and J). While, defects in lymph gland growth and progenitor homeostasis were evident in our previous study (Madhwal et al., 2020), the mechanism underlying the defects remained unaddressed. In this study, using the similar RNA*i* mediated genetic knock-down approach, we down-regulated each respective component of the GABA catabolic pathway in blood progenitor cells. Using progenitor specific drivers (*dome-MESO>GFP* and *TepIV>mCherry*) we assessed the role for this pathway in homeostatic conditions. We blocked: a) progenitor cell *Gat* function to perturb GABA uptake (Fig. 1C, J and Supp. Fig. 1A), b) *Ssadh* to perturb its breakdown into succinate (Fig. 1D, J and Supp. Fig. 1A, B), c) *sima* (Fig. 1E, J) and d) *Ldh* function to inhibit progenitor glycolysis (Fig. 1F, J). In all these conditions a significant reduction in lymph gland size was noticed. Blocking GABA uptake or the penultimate component of the pathway *Ldh*, in differentiating Hemolectin^+^ (Hml^+^) blood cells did not render any growth defect (Supp. Fig. 1 C, D, F and G). This implied a specific function for GABA breakdown in progenitor cells for lymph gland size control. Loss of *sima* from Hml^+^ differentiated blood cells (Supp. Fig. 1E and G) however led to a mild reduction in lymph gland size and indicated GABA independent functions of Sima in differentiating cells (Mukherjee et al., 2011). However, the requirement for Sima in progenitor cells for lymph gland growth was superior than observed in differentiating cells (Fig. 1E and J compared to Supp. Fig. 1E and G).

In progenitor cells, GABA inhibits Hph activity. This led us to ask its involvement in lymph gland growth. We found that over-expression of Hph in blood progenitor cells led to smaller sized lymph glands (Fig. 1G and K). This implied that Hph suppressed lymph gland growth. Blocking *Hph* function in *Gat^RNAi^* expressing progenitor cells (*Hph^RNAi^; Gat^RNAi^*) reverted the small lymph gland phenotype detected in *Gat^RNAi^* condition significantly (Fig. 1H and K). Loss of progenitor *Hph* function on its own, however, did not change lymph gland size (Fig. 1I and K), even though its gain of expression did (Fig. 1G and K). These data showed that, blood progenitor cells utilized GABA catabolic pathway to inhibit Hph function and this was necessary to promote lymph gland growth. Downstream of Hph, Sima protein and its metabolic target enzyme, Ldh, (Fig. 1A), when perturbed in the blood progenitor cells also led to small sized lymph glands (Fig. 1F, J). This showed their involvement as well in controlling growth of the hematopoietic tissue. Next, we investigated the mechanism by which GABA mediated Hph inhibition and downstream metabolic players controlled lymph gland growth.

### GABA catabolism in blood progenitors cells control their ROS levels

The progenitor cells of the lymph gland, exhibit a unique set of metabolic requirements, one of which is characterized by the presence of elevated levels of ROS in them (Fig. 2A). We observed that blocking *Gat* or *Ssadh* function in blood progenitor cells led to elevation of lymph gland ROS levels (Fig. 2A-C and F) which was detected utilizing dihydroethidium dye (DHE, Owusu-Ansah & Banerjee, 2009) (Refer to methods section for details). This suggested that intracellular GABA uptake and its breakdown in blood progenitor cells was necessary to maintain threshold levels of ROS in the lymph gland. The final metabolic output of GABA breakdown is succinate (Fig. 1A). Thus, when *Drosophila* larval mutants expressing *Gat^RNAi^* or *Ssadh^RNAi^* in blood progenitor cells were reared on food supplemented with succinate, this led to significant down-regulation of ROS levels (Fig. 2D-F). More importantly, succinate supplementation restored the associated lymph gland growth defect as well (Fig. 2D compared to B, E compared to C and G). These results suggested an underlying connection between GABA breakdown, ROS levels and lymph gland growth control.

**Figure 2.**
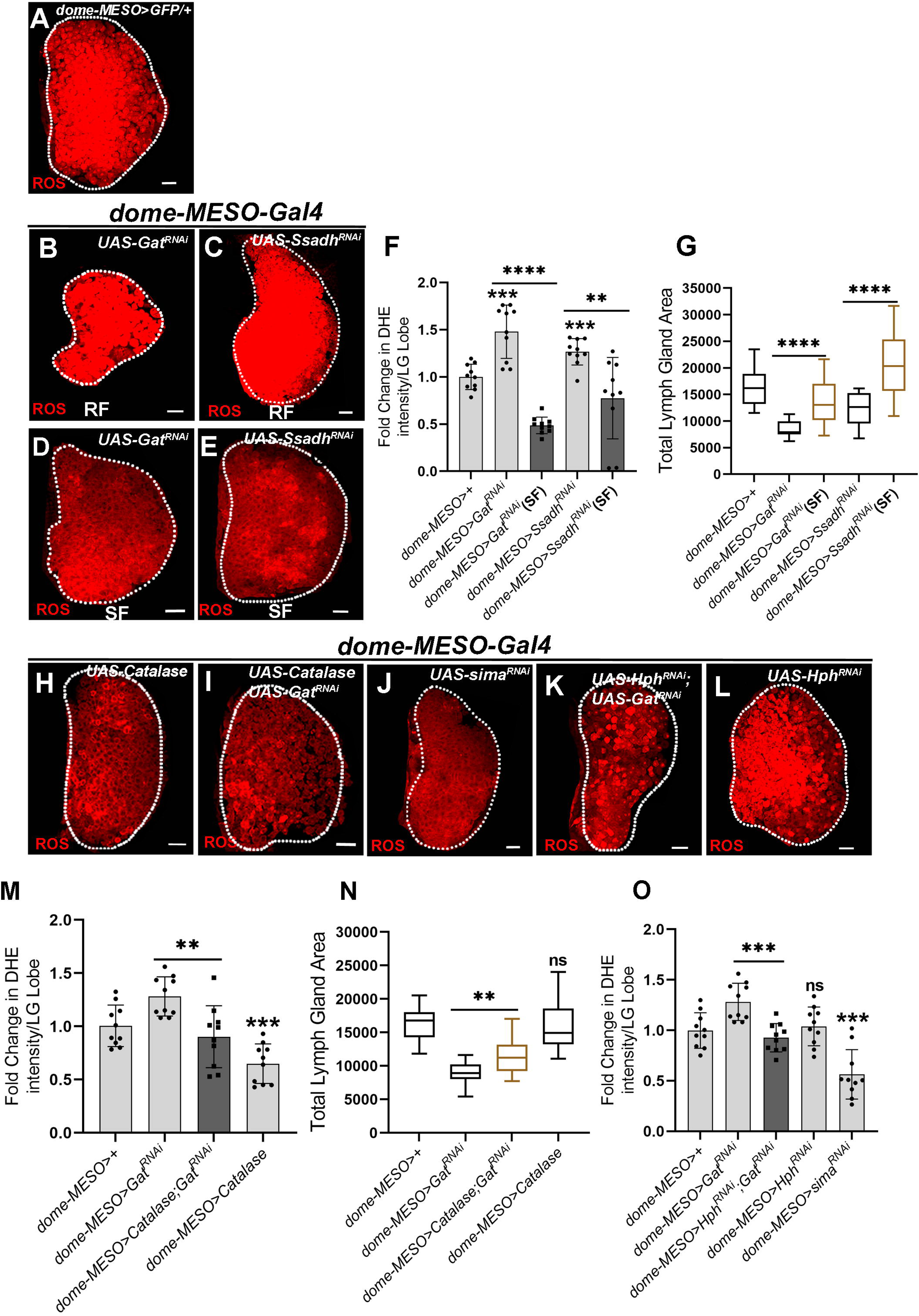
ROS regulation by GABA shunt pathway in *Drosophila* blood progenitors is important for lymph gland growth. DNA is marked with DAPI (blue). RF means Regular Food and SF means Succinate Food. Values are mean ± SD, asterisks mark statistically significant differences (*p<0.05; **p<0.01; ***p<0.001, ****p<0.0001, Student’s t-test for **F, M, O** and Mann-Whitney test for **G, N** is employed). Lymph glands are demarcated with white dotted line and are magnified for clarity of staining. Scale bar: 20µm. ‘n’ represents number of lymph gland lobes analysed. **(A)** Control (*dome-MESO>GFP/+*) lymph gland showing higher ROS levels in the blood progenitor cells, medullary zone (MZ). **(B, C)** Representative lymph gland images showing ROS levels in *Gat^RNAi^* and *Ssadh^RNAi^* on Regular Food (RF), **(B)** expressing *Gat^RNAi^* (*dome-MESO-Gal4, UAS-GFP; UAS-Gat^RNAi^*) and **(C)** expressing *Ssadh^RNAi^* (*dome-MESO-Gal4, UAS-GFP; UAS-Ssadh^RNAi^*) in blood progenitor cells leads to increase in ROS levels. For quantifications, refer to **F**. **(D, E)** Representative lymph gland images showing ROS levels in *Gat^RNAi^* and *Ssadh^RNAi^* on succinate food (SF), (**D)** succinate supplementation to *dome-MESO-Gal4, UAS-GFP; UAS-Gat^RNAi^* and (**E)** succinate supplementation to *dome-MESO-Gal4, UAS-GFP; UAS-Ssadh^RNAi^* rescues the increased ROS phenotype as compared to **(B)** and **(C)** respectively. For quantifications, refer to **F**. **(F)** Relative fold change in lymph gland ROS levels in *dome-MESO-Gal4, UAS-GFP*/+ (control, n=10), *dome-MESO-Gal4, UAS-GFP*; *UAS-Gat^RNAi^* (RF, n=10, p=0.0001), *dome-MESO-Gal4, UAS-GFP; UAS-Gat^RNAi^* (SF, n=10, p<0.0001), *dome-MESO-Gal4, UAS-GFP*; *UAS-Ssadh^RNAi^* (RF, n=10, p=0.0004) and *dome-MESO-Gal4, UAS-GFP; UAS-Ssadh^RNAi^* (SF, n=10, p=0.0030). **(G)** Quantifications of lymph gland area in *dome-MESO-Gal4, UAS-GFP*/+ (control, n=20), *dome-MESO-Gal4, UAS-GFP*; *UAS-Gat^RNAi^* (RF, n=20, p<0.0001), *dome-MESO-Gal4, UAS-GFP; UAS-Gat^RNAi^* (SF, n=20, p<0.0001), *dome-MESO-Gal4, UAS-GFP*; *UAS-Ssadh^RNAi^* (RF, n=20, p=0.0004) and *dome-MESO-Gal4, UAS-GFP; UAS-Ssadh^RNAi^* (SF, n=20, p<0.0001). **(H, I)** Representative lymph gland images showing ROS levels, **(H)** over-expressing *Catalase* (*dome-MESO-Gal4, UAS-GFP*; *UAS-Catalase*) shows reduction in ROS levels as compared to **(A)** control and **(I)** Over-expressing *Catalase* in *Gat^RNAi^* (*dome-MESO-Gal4, UAS-GFP*; *UAS-Catalase*; *UAS-Gat^RNAi^*) leads to rescue of lymph gland ROS defect as compared to **(B)** *Gat^RNAi^*. For quantifications, refer to **M**. **(J-L)** Representative lymph gland images showing ROS levels, (**J)** expressing *sima^RNAi^* in progenitor cells (*dome-MESO-Gal4, UAS-GFP; UAS-sima^RNAi^*) leads to reduction in ROS levels as compared to **(A)** control**, (K)** expressing *Hph^RNAi^* in *Gat^RNAi^* (*dome-MESO-Gal4, UAS-GFP*; *UAS-Hph^RNAi^*; *UAS-Gat^RNAi^*) leads to reduction in lymph gland ROS levels as compared to **(B)** *Gat^RNAi^* and (**L)** expressing *Hph^RNAi^* in progenitor cells (*dome-MESO-Gal4, UAS-GFP; UAS-Hph^RNAi^*) did not show any change in ROS levels as compared to **(A)** control. For quantifications, refer to **O. (M)** Relative fold change in lymph gland ROS levels in progenitor specific knock-down of *Gat*, over-expression of *Catalase* in *Gat^RNAi^* and over-expression of *Catalase*, *dome-MESO-Gal4, UAS-GFP/+* (control, n=10), *dome-MESO-Gal4, UAS-GFP; UAS-Gat^RNAi^* (n=10, p=0.0043), *dome-MESO-Gal4, UAS-GFP*; *UAS-Catalase*; *UAS-Gat^RNAi^* (n=10, p=0.0027), *dome-MESO-Gal4, UAS-GFP*; *UAS-Catalase* (n=10, p=0.0006). **(N)** Quantifications of lymph gland area in progenitor specific knock-down of *Gat*, over-expression of *Catalase* in *Gat^RNAi^* and over-expression of *Catalase*, *dome-MESO-Gal4, UAS-GFP/+* (control, n=20), *dome-MESO-Gal4, UAS-GFP; UAS-Gat^RNAi^* (n=20, p<0.0001), *dome-MESO-Gal4, UAS-GFP*; *UAS-Catalase*; *UAS-Gat^RNAi^* (n=20, p=0.0010), *dome-MESO-Gal4, UAS-GFP*; *UAS-Catalase* (n=20, p=0.4250). **(O)** Relative fold change in lymph gland ROS levels in progenitor specific knock-down of *Gat*, *Hph^RNAi^* in *Gat^RNAi^*, *sima*, and *Hph^RNAi^*. *dome-MESO-Gal4, UAS-GFP/+* (control, n=10), *dome-MESO-Gal4, UAS-GFP; UAS-Gat^RNAi^* (n=10, p=0.0024), *dome-MESO-Gal4, UAS-GFP; UAS-Hph^RNAi^*; *UAS-Gat^RNAi^* (n=10, p=0.0001), *dome-MESO-Gal4, UAS-GFP*; *UAS-Hph^RNAi^*; (n=10, p=0.6233) and *dome-MESO-Gal4, UAS-GFP; UAS-sima^RNAi^* (n=10, p=0.0002).

To address, if elevated ROS detected in GABA metabolic mutants was indeed the reason for lymph gland growth retardation, we asked if increasing progenitor ROS through independent means impacted lymph gland size. For this, we expressed RNA*i* against ROS scavenging enzyme, *superoxide dismutase 2 (sod2)* in progenitor cells. Similar to *Gat* and *Ssadh* loss of function, knockdown of *sod2* demonstrated an elevation in ROS (Supp. Fig. 2B and G) with a concomitant reduction in lymph gland size (Supp. Fig. 2H). This data implied that conditions leading to elevated ROS negatively affected lymph gland growth. Therefore, we undertook experiments to scavenge ROS in *Gat* and *Ssadh* mutant conditions and ask if this was sufficient to recover their growth defect. For this, we fed larvae expressing *Gat* and *Ssadh* RNA*i* in progenitor cells with N-acetylcysteine (NAC), a known antioxidant (Aldini et al., 2018; Niraula & Kim, 2019). This led to restoration of ROS levels in the mutant conditions (Supp. Fig. 2C-G) and significant recovery of lymph gland size (Supp. Fig. 2H). We undertook genetic means to scavenge ROS in *Gat^RNAi^* condition and this was undertaken by over-expressing ROS scavenging enzyme, catalase, in progenitor cells expressing *Gat^RNAi^*. Here as well, a significant reduction in ROS (Fig. 2H, I and M) and a significant recovery in lymph gland size (Fig. 2N and Supp. Fig. 2I-L) was evident. This suggested a role for blood progenitor ROS homeostasis via GABA metabolism in controlling lymph gland growth.

Down-stream of succinate, its inhibition of Hph activity and subsequent stabilization of progenitor Sima expression were demonstrated necessary for lymph gland growth (Fig. 1F, J). Hence, we explored the involvement of these components in progenitor ROS homeostasis. We tested Sima, as its loss led to small lymph glands, but interestingly, this condition did not recapitulate the increased ROS phenotype as seen with *Gat* or *Ssadh* loss of function. In fact a reduction in ROS levels was noted (Fig. 2J and O). However, loss of *Hph* in *Gat^RNAi^* expressing progenitor cells which restored growth, also restored ROS levels when compared to levels detected in *Gat^RNAi^* expressing cells (Fig. 2K and O). This implied modulating Hph function modulated ROS levels. However, any modulation of *Hph*, as seen with its loss in progenitor cells, did not manifest any change in ROS levels and remained comparable to controls (Fig. 2L and O). This suggested that succinate dependent regulation of progenitor *Hph* activity controlled lymph gland ROS independent of Sima and this function of Hph was sensitive to intracellular GABA. The independence of Hph function from Sima, also implied differential control of GABA in progenitor metabolism. We prospose that nhibition of Hph activity, by GABA limited excessive ROS levels in progenitor cells, whose accumulation retarded lymph gland growth. Secondly, Hph inhibition leading to Sima protein stability led to downstream Ldh function and positively controlled lymph gland growth. This dual control of GABA was also reflected in growth recovery brought about by loss of Hph function in *Gat^RNAi^* condition and succinate supplementation of *Gat^RNAi^* (SF) as opposed to growth recovery detected with ROS scavenging approaches. Feeding succinate or loss of *Hph* in *Gat^RNAi^* conditions showed superior growth recovery, while *Catalase* over-expression in *Gat^RNAi^* or *Gat^RNAi^* fed with NAC even though showed significant growth recovery, they were not comparable to the recovery detected with *Hph^RNAi^* or succinate (Supp. Fig. 2M). This corroborated with additional regulation by GABA on Sima and Ldh activity in coordinating lymph gland size, in addition to regulation of ROS.

### TCA activity, a prime producer of ROS in blood progenitor cells

Most studies in literature have implied Hph dependence on Hifα/Sima function to moderate ROS levels (Selak et al., 2005; Tannahill et al., 2013). The novel Sima independent link between GABA, Hph function and ROS regulation in lymph gland development led us to investigate the underlying mechanism.

In order to assess this control exerted by GABA, we first explored the source of developmental ROS in these cells. TCA cycle and its intermediates driving mitochondrial oxidative phosphorylation, is a significant center for ROS production (Mailloux et al., 2016; Quinlan et al., 2012; Starkov et al., 2004; Woolbright et al., 2019). Thus, we asked if TCA activity in lymph gland progenitor cells was responsible for ROS detected in them. For this, as a proxy for measuring TCA activity, we assessed expression levels of key TCA enzymes PDK and PDH, using antibody that detect their active and inactive forms. PDH enzyme converts pyruvate to acetyl-CoA and drives the TCA cycle (Wang et al., 2016). PDH activity is regulated at the level of its phosphorylation, where the phosphorylated form (pPDH) marks inactive enzymatic state and represses TCA activity. Phosphorylation of PDH is mediated by PDK, whose phosphorylated form (pPDK), marks active PDK state and hence decreased TCA activity (Fig. 3A).

**Figure 3.**
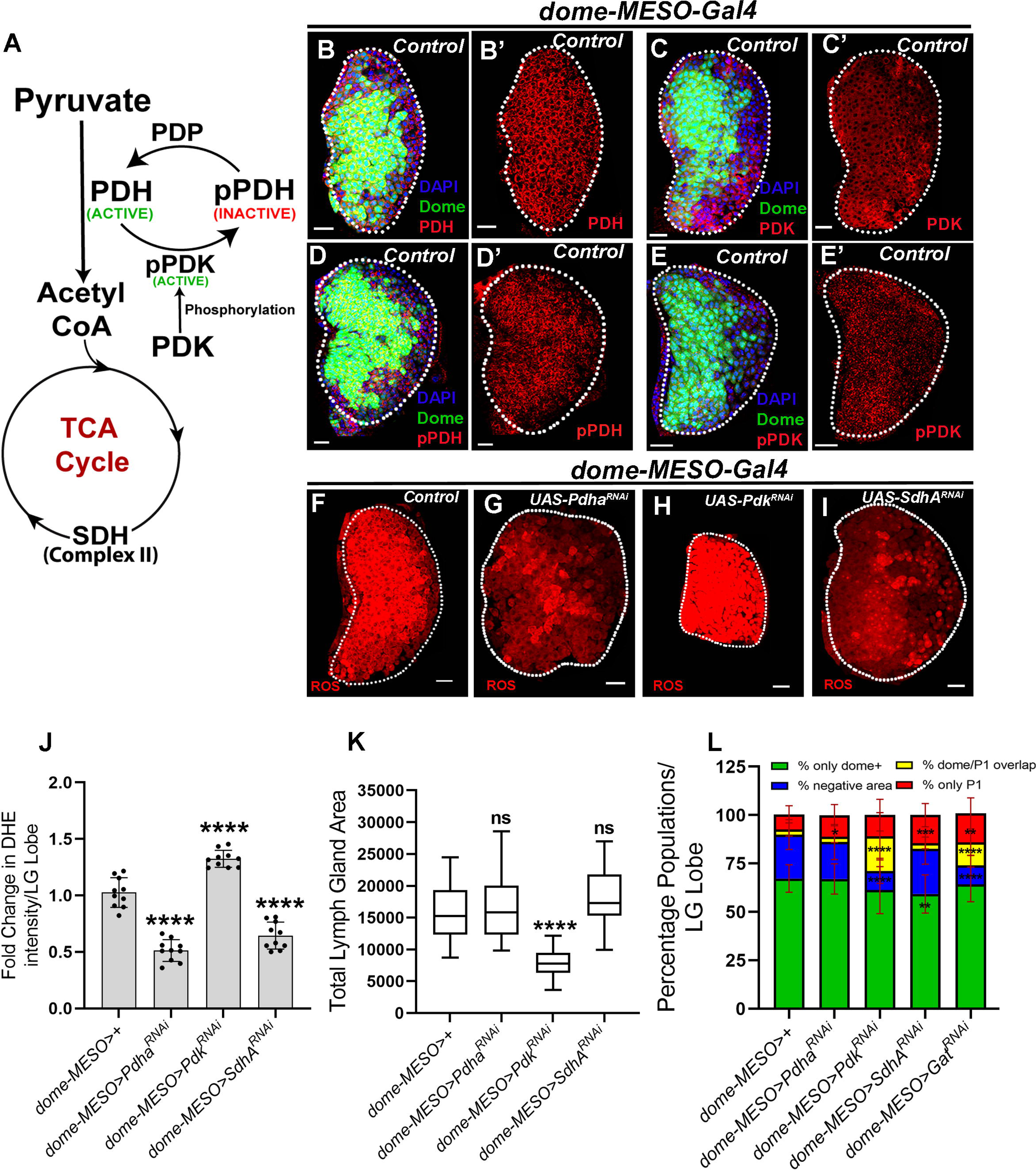
TCA cycle contributes to Medullary zone ROS and regulates lymph gland growth. DNA is marked with DAPI (blue). dome^+^ (green) marks the lymph gland blood progenitor cells (medullary zone, MZ) and dome^-^ region marks the cortical zone (CZ). Values are mean ± SD, asterisks mark statistically significant differences (*p<0.05; **p<0.01; ***p<0.001, ****p<0.0001, Student’s t-test for **J, L** and Mann-Whitney test for **K** is employed). Lymph glands are demarcated with white dotted line. Scale bar: 20µm. ‘n’ represents number of lymph gland lobes analysed. **(A)** Schematic representation showing regulation of pyruvate entry and conversion to acetyl-CoA by PDH enzyme (active). PDH enzyme is phosphorylated by PDK which makes it inactive and PDP dephosphorylates pPDH (inactive) to PDH (active). Pyruvate converts into acetyl-CoA which fuels into the TCA cycle. **(B-C’)** Representative lymph gland images showing PDH (red) and PDK1 (red) in control lymph gland (*dome-MESO>GFP/+*), **(B,B’)** PDH expression in the *dome-MESO>GFP/+* **(B)** with dome+ overlap (green) and **(B’)** without dome+ overlap, **(C,C’)** PDK1 expression in the *dome-MESO>GFP/+* **(C)** with dome+ overlap (green) and **(C’)** without dome+ overlap, show no alteration in the PDH and PDK1 levels in the dome^+^ (MZ) and dome^-^ region (CZ). **(D-E’)** Representative lymph gland images showing pPDH (red) and pPDK (red) in control lymph gland (*dome-MESO>GFP/+*), **(D,D’)** pPDH expression in the *dome-MESO>GFP/+* **(D)** with dome+ overlap (green) and **(D’)** without dome+ overlap, **(E, E’)** pPDK expression in the *dome-MESO>GFP/+* **(E)** with dome+ overlap (green) **(E’)** and without dome+ overlap, show more pPDH and pPDK levels in the dome^+^ (MZ) as compared to dome^-^ region (CZ). **(F-I)** Representative lymph gland images showing ROS levels, compared to **(F)** control (*dome-MESO-Gal4, UAS-GFP*/+); **(G)** expressing *Pdha^RNAi^* (*dome-MESO-Gal4, UAS-GFP*; *UAS-Pdha^RNAi^*) leads to reduction in ROS levels, **(H)** expressing *Pdk^RNAi^* (*dome-MESO-Gal4, UAS-GFP*; *UAS-Pdk^RNAi^*) elevates ROS levels in lymph gland blood progenitors and **(I)** expressing *SdhA^RNAi^* (*dome-MESO-Gal4, UAS-GFP*; *UAS-SdhA^RNAi^*) leads to reduction in ROS levels. For quantifications, refer to **J**. **(J)** Relative fold change in lymph gland ROS levels in *dome-MESO-Gal4, UAS-GFP*/+ (control, n=10), *dome-MESO-Gal4, UAS-GFP*; *UAS-Pdha^RNAi^* (n=10, p<0.0001), *dome-MESO-Gal4, UAS-GFP*; *UAS-Pdk^RNAi^* (n=10, p<0.0001) and *dome-MESO-Gal4, UAS-GFP*; *UAS-SdhA^RNAi^* (n=10, p<0.0001). **(K)** Quantifications of lymph gland size in *dome-MESO-Gal4, UAS-GFP*/+ (control, n=20), *dome-MESO-Gal4, UAS-GFP*; *UAS-Pdha^RNAi^* (n=20, p=0.8201), *dome-MESO-Gal4, UAS-GFP*; *UAS-Pdk^RNAi^* (n=20, p<0.0001) and *dome-MESO-Gal4, UAS-GFP*; *UAS-SdhA^RNAi^* (n=20, p=0.0596). **(L)** Quantifications of lymph gland differentiation status shown as percentage of only dome^+^(green), dome^-^P1^-^(blue), dome^+^P1^+^(yellow) and only P1^+^(red) populations per lymph gland lobe. p-values are presented in the preceding order. *dome-MESO-Gal4, UAS-GFP*/+ (control, n=20), *dome-MESO-Gal4, UAS-GFP*; *UAS-Pdha^RNAi^* (n=20, p=0.9445, 0.1749, 0.9926, 0.0337), *dome-MESO-Gal4, UAS-GFP*; *UAS-Pdk^RNAi^* (n=20, p=0.0663, <0.0001, <0.0001, 0.1328), *dome-MESO-Gal4, UAS-GFP*; *UAS-SdhA^RNAi^* (n=20, p=0.0058, 0.7883, 0.7997, 0.0003) and *dome-MESO-Gal4, UAS-GFP*; *UAS-Gat^RNAi^* (n=20, p=0.2432, <0.0001, <0.0001, 0.0012). Representative lymph gland images are shown in **Supplementary** Fig. 3G**-J’’**.

Immuno-histochemical analysis of third instar larval lymph glands against PDH^total^, PDK^total^, active pPDK, and inactive pPDH was undertaken. PDH^total^ and PDK^total^ showed uniform expression in all cells of a 3^rd^ instar larval lymph gland (Fig. 3B-C’). Compared to this, levels of pPDH (Fig. 3D, D’) and pPDK (Fig. 3E, E’) was specifically elevated in dome^+^ progenitor cells when compared to their levels detected in dome^-^ differentiating cells (Fig. 3D-E’).

These expression data suggested that in homeostasis, blood progenitor cells maintained substantial fraction of PDH in an inactive state (pPDH). The increased threshold of active PDK (pPDK) in progenitor cells highlighted PDK’s regulation to limit PDH function and suggested moderation of TCA activity in these cells.

We employed genetic means to perturb *Pdh* and *Pdk* enzymes by employing RNA*i* against them and assessed changes in progenitor ROS generation and consequently blood progenitor development. First, we confirmed the specificity of the RNA*i* lines by undertaking expression analysis of pPDH and PDH^total^ antibodies in lymph glands expressing *Pdha^RNAi^* and *Pdk^RNAi^* (Supp. Fig 3). A striking down-regulation of PDH^total^ (Supp. Fig 3A, B and K) and pPDH (Supp. Fig 3D, E and L) expression was seen in *Pdha^RNAi^* expressing lymph glands. In *Pdk^RNAi^* expressing lymph glands, PDH^total^ expression remained unaffected (Supp. Fig 3C, K), but the expression of phosphorylated form (pPDH) was reduced (Supp. Fig 3F, L). Together, these data confirmed the specificity of the RNA*i* lines. The specific loss of pPDH in *Pdk^RNAi^* condition also showed that the phosphorylation of PDH in progenitor cells was reliant on Pdk function.

RNA*i* mediated knock-down of *Pdha* expression in blood progenitor cells, led to a 50% reduction in ROS levels when compared to levels detected in control lymph glands (Fig. 3G compared to F and J). Importantly, loss of progenitor *Pdha* expression did not impede lymph gland growth and they remained comparable to control conditions (Fig. 3G, K and Supp. Fig. 3G, H). On the other hand, blocking *Pdk* expression in blood progenitor cells, led to almost 1.5-fold increase in ROS levels (Fig. 3H compared to F and J). Here, a dramatic reduction in lymph gland size was noticed (Fig. 3H, K and Supp. Fig. 3I). This was further confirmed by using another independent RNA*i* against *Pdk*, which phenocopied the small lymph gland size (Supp. Fig. 3M) as detected with loss of *Gat*, *Ssadh* or *sod2* (Supp. Fig. 2H) conditions. Thus, taken together, *Pdha* loss-of-function data implied PDH function in facilitating production of progenitor ROS and the *Pdk* loss-of-function data implied PDK activity in controlling progenitor ROS levels and lymph gland size.

Consistent with TCA involvement in OXPHOS, progenitor cell specific down-regulation of TCA cycle enzyme, *Succinate dehydrogenase A (SdhA)*, also a component of Complex II of the mitochondrial electron transport chain (ETC), lowered progenitor ROS levels (Fig. 3I, J). In this genetic condition, lymph gland sizes remained unaffected (Fig. 3H, K and Supp. Fig. 3J) as well. These loss of function data were consistent with phenotypes detected with *Pdh* loss in progenitor cells, that functioned upstream in the TCA cycle. The data showed that in homeostatic conditions, PDH dependent entry of pyruvate into the TCA and the subsequent oxidation and activation of mitochondrial ETC via SDH caused ROS production in blood progenitor cells. Lowering ROS and TCA function was however not necessary for lymph gland growth, but the regulatory step via Pdk, implicated in moderating PDH activity to limit excessive TCA activity. This was necessary to limit precocious ROS generation in progenitor cells and support lymph gland growth.

If modulation of the TCA enzymes also affected progenitor homeostasis was examined. This was addressed by staining lymph glands for differentiation marker P1 along with assessment of progenitor population undertaken by analysing dome-GFP reporter expression in the above-mentioned genetic conditions. In control lymph glands, we observed that 60-70% area of lymph gland was Dome^+^, 15-20% area was dome^-^ but P1^+^ and remaining 20-25% was negative for both the markers (Fig. 3L and Supp. Fig. 3 G-G”). Loss of *Pdha* function (low TCA) from progenitor cells, did not alter progenitor homeostasis and remained comparable to controls. A mild increase in plasmatocytes population was observed (Fig. 3L and Supp. Fig. 3 H-H”). Like *Pdha* loss of function, *SdhA^RNAi^* also did not show any striking change in progenitor status but here as well the mild increase in the plasmatocytes population along with slight decrease in dome^+^ area was apparent (Fig. 3L and Supp. Fig. 3 J-J”). This implied a requirement for TCA in differentiation. In *Pdk^RNAi^* condition, however, changes were more dramatic. An increase in dome^+^ population was apparent, along with this, an increase in P1 population (Fig. 3L and Supp. Fig. 3I-I”) and reduction in dome^-^P1^-^ population was detected. Importantly, in *Pdk^RNAi^* condition, as opposed to *Pdha^RNAi^* or *SdhA^RNAi^,* we observed a subset of dome^+^ cells overlapping with P1^+^ cells (Supp. Fig. 3I). This population was undetectable in homeostasis and their appearance in *Pdk^RNAi^* implied a role for Pdk function in maintenance of progenitor cell state and their proper differentiation. Given the similarities between *Pdk* and GABA-metabolic mutants, we assessed dome^+^ and P1^+^ cells in *Gat^RNAi^* mutant condition. Here as well, we observed a similar trend and identified dome^+^P1^+^ overlapping cells (Fig. 3L). These progenitor changes detected in *Pdk ^RNAi^* and *Gat^RNAi^* conditions implied that in addition to controlling lymph gland size, moderating TCA activity via GABA was important in maintaining differentiation of progenitor cells.

### GABA-catabolism regulates TCA activity by regulating PDK function and moderates ROS generation in blood progenitor cells

Based on the phenotypic similarities observed between GABA and *Pdk* loss of function data, we hypothesized an increase in TCA activity in GABA catabolic mutants, that led to the small lymph gland and differentiation phenotypes. We investigated the levels of PDH^total^, PDK^total^, active PDK (phospho-PDK), inactive form of PDH (p-PDH) in GABA metabolic pathway mutants (Fig. 4 and Supp. Fig. 4). Expression levels of PDH^total^, PDK^total^ remained unchanged upon loss of progenitor *Gat* and *Ssadh* expression (Fig. 4A-F and Supp. Fig. 4A, B). This showed that changes in GABA metabolism did not alter the production of these enzymes. However, we observed specific down-regulation in pPDH (inactive PDH, Fig. 4G-I, and Supp. Fig. 4C) and pPDK (active PDK, Fig. 4J-L and Supp. Fig. 4D) expression upon loss of *Gat* and *Ssadh* expression in progenitor cells. The specific reduction in pPDH expression, implied increased fraction of active PDH enzyme and enhanced TCA activity in GABA metabolic mutants.

**Figure 4.**
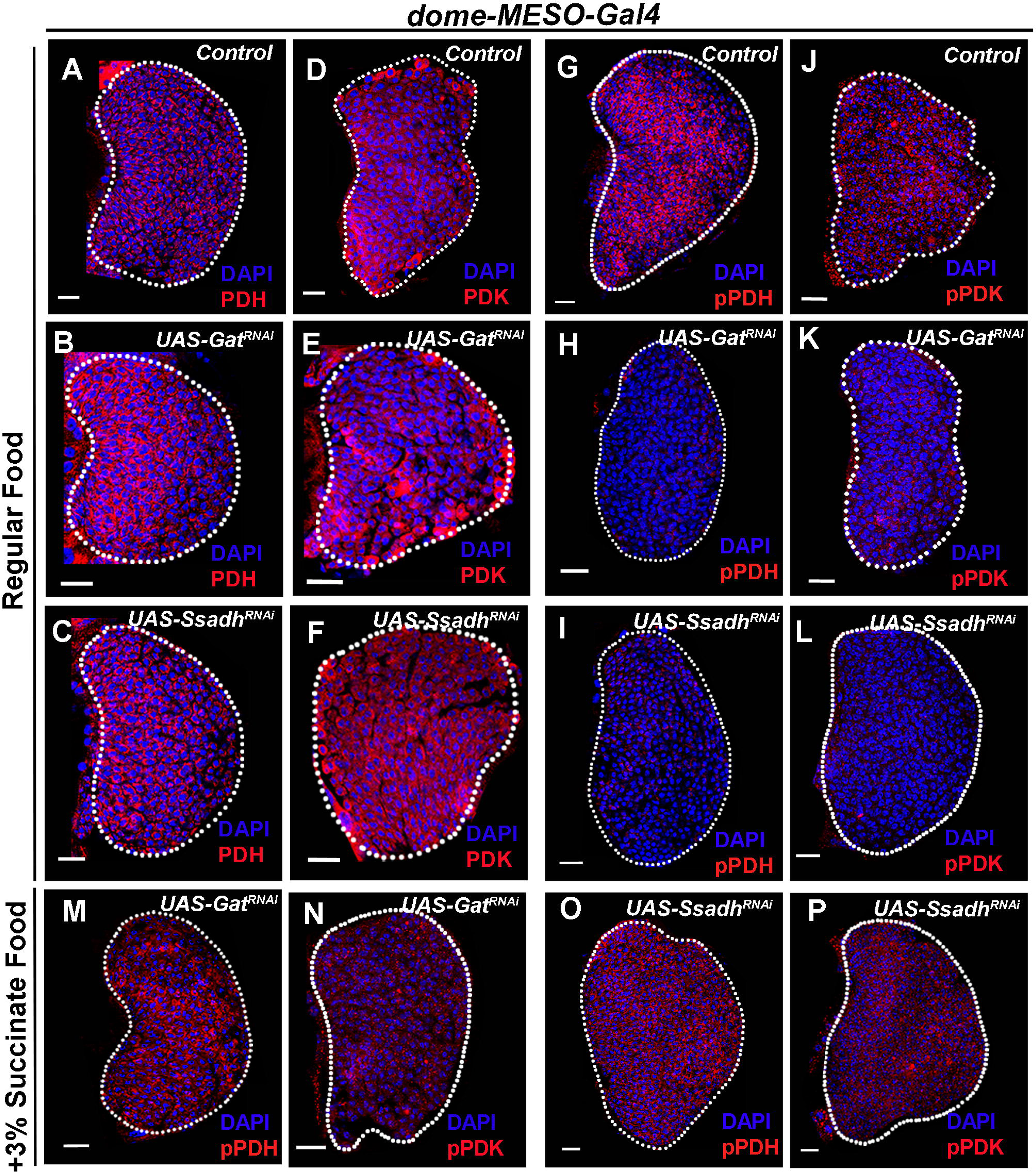
GABA catabolism controls PDH activity to regulate lymph gland growth. DNA is marked with DAPI (blue). RF means Regular Food and SF means Succinate Food. Lymph glands are demarcated with white dotted line and are magnified for clarity of staining. Scale bar: 20µm. **(A-C)** Representative lymph gland images showing the expression PDH in progenitor specific knock-down of *Gat* and *Ssadh*, **(A)** control (*dome-MESO>GFP/+*) showing PDH expression**, (B)** expressing *Gat^RNAi^* in progenitor cells (*dome-MESO-Gal4, UAS-GFP; UAS-Gat^RNAi^*) and **(C**) expressing *Ssadh^RNAi^* in progenitor cells (*dome-MESO-Gal4, UAS-GFP; UAS-Ssadh^RNAi^*) does not show reduction in PDH levels in the medullary zone as compared to **(A)** control (*dome-MESO>GFP/+*). For quantifications, refer to **Supplementary** Fig. 4A. **(D-F)** Representative lymph gland images showing the expression PDK1 in progenitor specific knock-down of *Gat* and *Ssadh*, **(D)** control (*dome-MESO>GFP/+*) showing PDK1 expression**, (E)** expressing *Gat^RNAi^* in progenitor cells (*dome-MESO-Gal4, UAS-GFP; UAS-Gat^RNAi^*) and **(F**) expressing *Ssadh^RNAi^* in progenitor cells (*dome-MESO-Gal4, UAS-GFP; UAS-Ssadh^RNAi^*) does not show reduction in PDK1 levels in the medullary zone as compared to **(D)** control (*dome-MESO>GFP/+*). For quantifications, refer to **Supplementary** Fig. 4B. **(G-I)** Representative lymph gland images showing the expression pPDH in progenitor specific knock-down of *Gat* and *Ssadh*, **(G)** control (*dome-MESO>GFP/+*) showing pPDH expression**, (H)** expressing *Gat^RNAi^* in progenitor cells (*dome-MESO-Gal4, UAS-GFP; UAS-Gat^RNAi^*) and **(I**) expressing *Ssadh^RNAi^* in progenitor cells (*dome-MESO-Gal4, UAS-GFP; UAS-Ssadh^RNAi^*) leads to reduction in pPDH levels in the medullary zone as compared to **(G)** control (*dome-MESO>GFP/+*). For quantifications, refer to **Supplementary** Fig. 4C. **(J-L)** Representative lymph gland images showing the expression pPDK in progenitor specific knock-down of *Gat* and *Ssadh*, **(J)** control (*dome-MESO>GFP/+*) showing pPDK expression**, (K)** expressing *Gat^RNAi^* in progenitor cells (*dome-MESO-Gal4, UAS-GFP; UAS-Gat^RNAi^*) and **(L**) expressing *Ssadh^RNAi^* in progenitor cells (*dome-MESO-Gal4, UAS-GFP; UAS-Ssadh^RNAi^*) does not show reduction in pPDK levels in the medullary zone as compared to **(J)** control (*dome-MESO>GFP/+*). For quantifications, refer to **Supplementary** Fig. 4D. **(M-P)** Succinate supplementation into progenitor specific loss of *Gat* and *Ssadh* lymph glands rescues the pPDH and pPDK phenotypes. (**M, N)** succinate supplementation in *dome-MESO-Gal4, UAS-GFP; UAS-Gat^RNAi^* rescues the **(M)** pPDH and **(N)** pPDK levels as compared to **(H, K)** *dome-MESO-Gal4, UAS-GFP; UAS-Gat^RNAi^* on RF and **(O, P)** succinate supplementation to *dome-MESO-Gal4, UAS-GFP; UAS-Ssadh^RNAi^* rescues the **(O)** pPDH and **(P)** pPDK levels as compared to as compared to **(I, L)** *dome-MESO-Gal4, UAS-GFP; UAS-Ssadh^RNAi^* on RF, respectively. For quantifications, refer to **Supplementary** Fig. 4C**, D**.

The specific reduction in pPDK levels, suggested GABA function in maintaining active PDK state in progenitor cells to limit PDH activity and suppress TCA. We therefore investigated this by down-regulating components of the TCA cycle in *Gat^RNAi^* condition and assessed for changes in lymph gland ROS and growth phenotypes. Interestingly, down-regulation of *Pdha* or *SdhA* expression in *Gat^RNAi^* expressing progenitor cells corrected lymph gland elevated ROS phenotype almost comparable to levels detected in controls (Fig. 5A-E). In addition to this, growth defect in *Gat^RNAi^* condition was restored significantly (Fig. 5F-J). *SdhA^RNAi^; Gat^RNAi^* combination recovered lymph gland size almost comparable to controls (Fig. 5J) and suggested that its activity in GABA metabolic mutants was the key source of precocious ROS generation leading to the growth defect. We also assessed for differentiation status and observed recovery in progenitor differentiation status in *Gat^RNAi^* condition upon loss of *PdhA* and *SdhA* function in these cells. Dome^+^P1^+^ double positive cells that were detected in *Gat^RNAi^* condition were no longer detectable in *Pdha^RNAi^; Gat^RNAi^* or in *SdhA^RNAi^; Gat^RNAi^* condition (Fig. 5F-I’’ and K). Taken together, these data suggested that GABA function in progenitor cells via regulating PDK activity, controlled PDH dependent pyruvate entry into TCA and OXPHOS and further controlled precocious ROS generation. This regulation supported growth and proper differentiation of progenitor cells.

**Figure 5.**
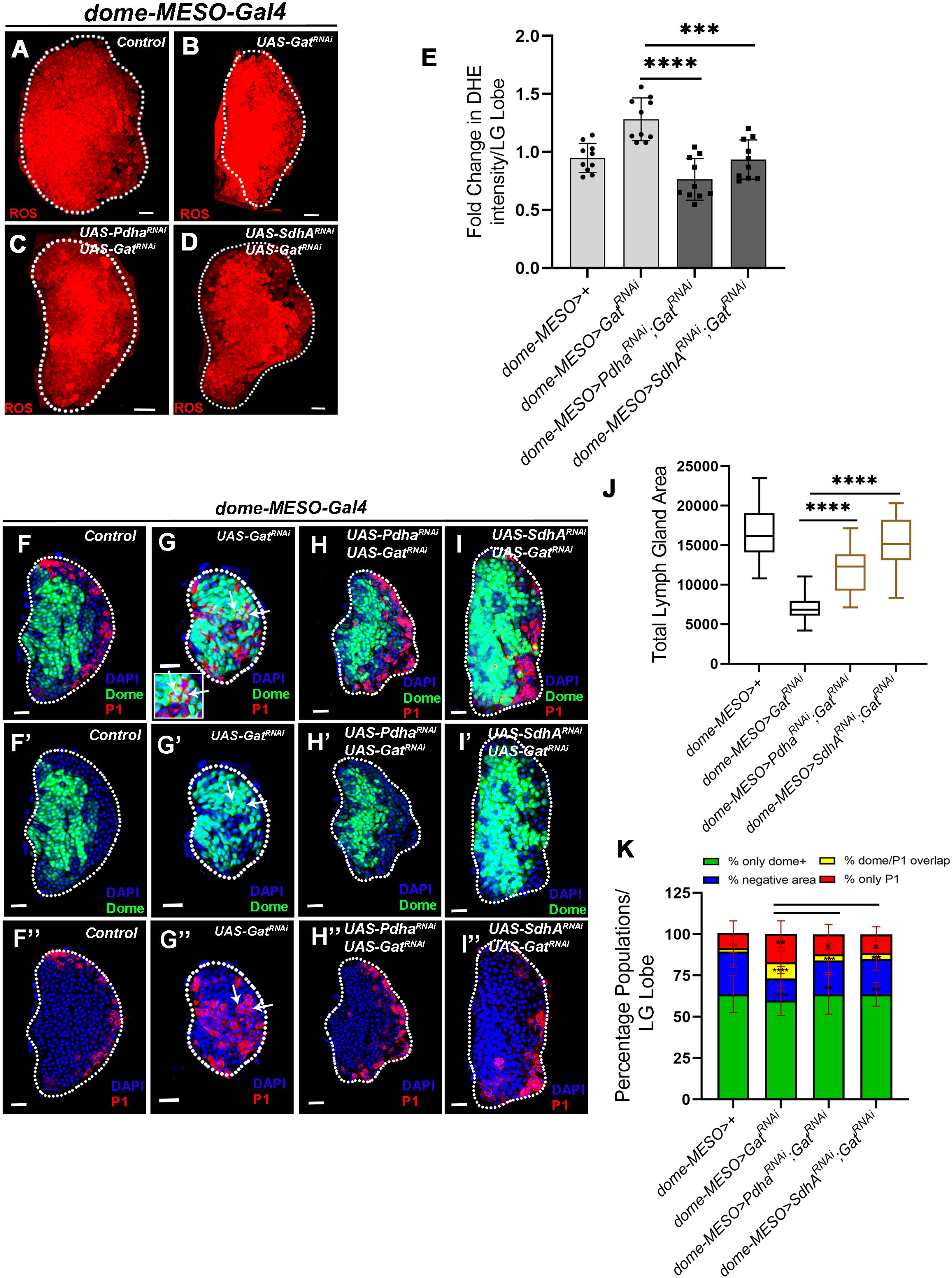
GABA catabolism regulates TCA activity to regulate ROS and lymph gland growth. DNA is marked with DAPI (blue). Values are mean ± SD, asterisks mark statistically significant differences (*p<0.05; **p<0.01; ***p<0.001, ****p<0.0001, Student’s t-test for **E, K** and Mann-Whitney test for **J** is employed). Lymph glands are demarcated with white dotted line and are magnified for clarity of staining. White arrows in **G-G”** indicates the dome^+^P1^+^ overlapping cells. Scale bar: 20µm. ‘n’ represents number of lymph gland lobes analysed. **(A-D)** Representative lymph gland images showing ROS levels in the **(A)** control (*dome-MESO-Gal4, UAS-GFP*/+), **(B)** expressing *Gat^RNAi^* in progenitor cells (*dome-MESO-Gal4, UAS-GFP; UAS-Gat^RNAi^*) leads to elevation in ROS levels, **(C**) expressing *Pdha^RNAi^* in *Gat^RNAi^* (*dome-MESO-Gal4, UAS-GFP; UAS-Pdha^RNAi^; UAS-Gat^RNAi^*) and **(D**) expressing *SdhA^RNAi^* in *Gat^RNAi^* (*dome-MESO-Gal4, UAS-GFP; UAS-SdhA^RNAi^; UAS-Gat^RNAi^*) rescues the increased ROS phenotype of **(B)** *Gat^RNAi^*. For quantifications refer to **E**. **(E)** Relative fold change in lymph gland ROS levels in *dome-MESO-Gal4, UAS-GFP*/+ (control, n=10), *dome-MESO-Gal4, UAS-GFP*; *UAS-Gat^RNAi^* (n=10, p=0.0002), *dome-MESO-Gal4, UAS-GFP; UAS-Pdha^RNAi^; UAS-Gat^RNAi^* (n=10, p<0.0001), and *dome-MESO-Gal4, UAS-GFP*; *UAS-SdhA^RNAi^; UAS-Gat^RNAi^* (n=10, p=0.0004). **(F-I’’)** Representative images showing lymph gland growth and differentiation status, **(F-F’’)** control (*dome-MESO-Gal4, UAS-GFP*/+) lymph gland showing **(F, F’)** dome^+^ (green), **(F)** dome^-^P1^-^(blue, DAPI) and **(F, F’’)** P1^+^ (red) population, **(G-G’’)** expressing *Gat^RNAi^* in progenitor cells (*dome-MESO-Gal4, UAS-GFP; UAS-Gat^RNAi^*) leads to small lymph gland size and appearance of **(G)** dome^+^p1^+^ overlap population along with an increase in **(G”)** P1 population, **(H-H’’**) expressing *Pdha^RNAi^* in *Gat^RNAi^* (*dome-MESO-Gal4, UAS-GFP; UAS-Pdha^RNAi^; UAS-Gat^RNAi^*) and **(I-I’’**) expressing *SdhA^RNAi^* in *Gat^RNAi^* (*dome-MESO-Gal4, UAS-GFP; UAS-SdhA^RNAi^; UAS-Gat^RNAi^*) rescues the lymph gland growth and differentiation defect of **(G-G’’)** *Gat^RNAi^*. For quantifications refer to **J and K**. **(J)** Quantifications of lymph gland size in *dome-MESO-Gal4, UAS-GFP*/+ (control, n=20), *dome-MESO-Gal4, UAS-GFP*; *UAS-Gat^RNAi^* (n=20, p<0.0001), *dome-MESO-Gal4, UAS-GFP; UAS-Pdha^RNAi^; UAS-Gat^RNAi^* (n=20, p<0.0001), and *dome-MESO-Gal4, UAS-GFP*; *UAS-SdhA^RNAi^; UAS-Gat^RNAi^* (n=20, p<0.0001). **(K)** Quantifications of lymph gland differentiation status shown as percentage of only dome^+^(green), dome^-^P1^-^(blue), dome^+^P1^+^(yellow) and only P1^+^(red) populations per lymph gland lobe. p-values are presented in the preceding order. *dome-MESO-Gal4, UAS-GFP*/+ (control, n=20) and *dome-MESO-Gal4, UAS-GFP*; *UAS-Gat^RNAi^* (n=20, p=0.2558, <0.0001, <0.0001, 0.0022 in comparison to control), *dome-MESO-Gal4, UAS-GFP; UAS-Pdha^RNAi^; UAS-Gat^RNAi^* (n=20, p=0.2918, 0.0032, 0.0006, 0.0312 in comparison to *Gat^RNAi^*), and *dome-MESO-Gal4, UAS-GFP*; *UAS-SdhA^RNAi^; UAS-Gat^RNAi^* (n=17, p=0.1891, 0.0014, 0.0015, 0.0128 in comparison to *Gat^RNAi^*).

### GABA-catabolism via Hph independent of Sima controls ROS homeostasis in blood progenitor cells

The mechanism by which GABA catabolic pathway regulated pPDK levels was explored. The restoration of both ROS and lymph gland growth defect with succinate supplemented diet implied regulation of TCA activity via succinate (Fig. 2A-G). Thus, we examined expression levels of PDH and PDK in succinate supplemented conditions. Compared to pPDH and pPDK levels detected in *Gat* and *Ssadh* RNA*i* expressing animals raised on regular food (Fig. 4M-P and Supp. Fig. 4C, D), we observed that succinate supplemented diet restored levels of pPDK (Fig. 4N, P compared to K, L respectively) and consequently pPDH (Fig. 4M, O compared to H, I respectively) to almost levels detected in wild-type control. Since GABA derived succinate functioned by inhibiting Hph activity in progenitor cells, expression levels of pPDH and pPDK in *Hph^RNAi^*; *Gat^RNAi^* expressing animals was assessed. Here as well, a significant recovery of pPDH and pPDK levels was noted (Fig. 6E, F compared to C, D and Supp Fig. 4E, F). This implied that loss of *Hph* function in *Gat^RNAi^* expressing progenitor cells was sufficient to recover pPDK and consequently pPDH levels. Inhibition of Hph in progenitors cells independently led to a significant upregulation in pPDH and pPDK levels (Fig. 6G, H compared to A, B and Supp. Fig. 4E, F) and implied Hph function in inhibiting PDK activity. Even though, *Hph* loss elevated PDK function, it did not alter lymph gland ROS (Fig. 2L) or its growth (Fig. 1I). These data suggested modulation of PDK leading to ROS by Hph was sensitive to levels of intracellular GABA. This was reflected in *Hph^RNAi^*; *Gat^RNAi^* expressing progenitor cells, where loss of Hph led to recovery of PDK activity, restoration of PDH suppression, lowering of TCA and consequently ROS levels.

**Figure 6.**
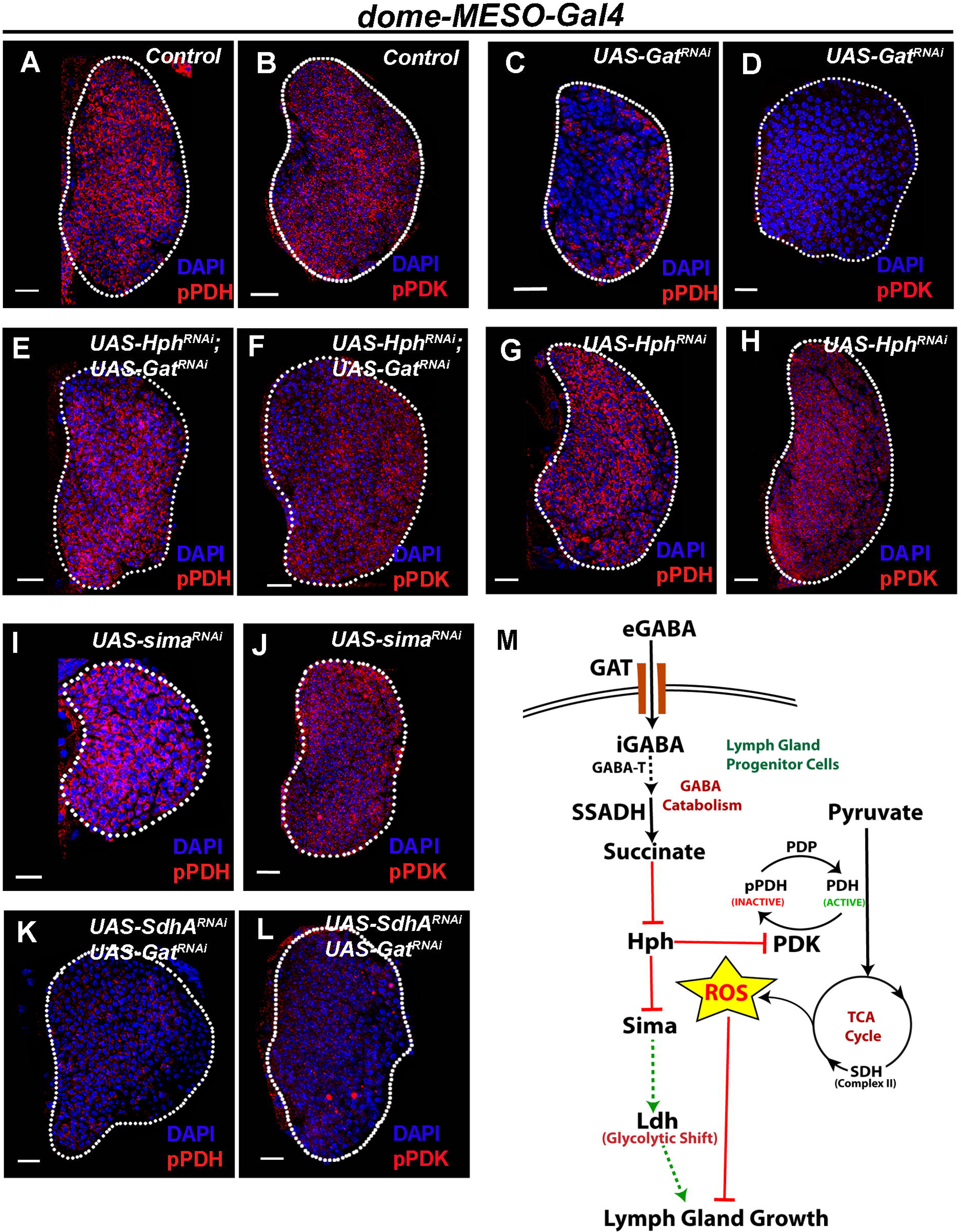
GABA catabolism regulates Hph activity to control lymph gland growth and regulate ROS homeostasis. DNA is marked with DAPI (blue). Lymph glands are demarcated with white dotted line and are magnified for clarity of staining. Scale bar: 20µm. **(A-L)** Representative lymph gland images showing the expression of pPDH and pPDK, **(A, B)** control (*dome-MESO>GFP/+*) showing **(A)** pPDH and **(B)** pPDK expression**, (C, D)** expressing *Gat^RNAi^* in progenitor cells (*dome-MESO-Gal4, UAS-GFP; UAS-Gat^RNAi^*) show reduction in **(C)** pPDH and **(D)** pPDK expression in the medullary zone as compared to **(A,B)** control (*dome-MESO>GFP/+*), **(E, F)** Expressing *Hph^RNAi^* in *Gat^RNAi^* (*dome-MESO-Gal4, UAS-GFP*; *UAS-Hph^RNAi^*; *UAS-Gat^RNAi^*) leads to recovery of **(E)** pPDH and **(F)** pPDK levels in lymph gland blood progenitors as compared to **(C, D)** *Gat^RNAi^***, (G, H)** expressing *Hph^RNAi^* (*dome-MESO-Gal4, UAS-GFP*; *UAS-Hph^RNAi^*) shows increase in **(G)** pPDH and **(H)** pPDK levels in blood progenitor cells as compared to **(A, B)** control, **(I, J)** expressing *sima^RNAi^* (*dome-MESO-Gal4, UAS-GFP*; *UAS-sima^RNAi^*) in blood progenitor cells does not show any change in **(I)** pPDH and **(J)** pPDK levels in blood progenitor cells as compared to **(A, B)** control, *dome-MESO-Gal4, UAS-GFP*/+, **(K, L)** Expressing *SdhA^RNAi^* in *Gat^RNAi^* (*dome-MESO-Gal4, UAS-GFP*; *UAS-SdhA^RNAi^*; *UAS-Gat^RNAi^*) does not show recovery of **(K)** pPDH and **(L)** pPDK levels in lymph gland blood progenitors as compared to **(C, D)** *Gat^RNAi^*. For quantifications refer to **Supplementary** Fig. 4E**, F**. **(M) Schematic representation showing the mechanism of GABA catabolic pathway in regulating lymph gland growth.** GABA catabolism derived succinate mediates inhibition of Hph enzyme activity which is essential for maintenance of an active phosphorylated form of PDK (pPDK) in blood progenitor cells. PDK phosphorylates PDH (pPDH) and inactivates it, which leads to lowering of TCA activity and subsequent ROS generation in progenitor cells. Elevated ROS in these cells results in the reduction of the overall lymph gland size and aberrant differentiation. In addition to this, GABA dependent regulation of Hph activity stabilizes Sima, which independent of ROS, positively controls lymph gland growth by mediating the glycolytic shift via Ldh, that metabolically programmes progenitor cells to support growth.

Hph function in progenitor cells in the context of ROS regulation was independent of Sima. This notion was supported by PDH and PDK expression levels in *sima^RNAi^* expressing progenitor cells whose levels did not change (Fig. 6I, J compared to A, B and Supp. Fig. 4E, F). This corroborated with the independence of ROS detected in Sima condition as well. These data reinforced alternative mechanisms employed by Sima to coordinate lymph gland growth. Based on involvement of Ldh in regulating lymph gland growth as demonstrated in the previous section, we propose Sima function in activating Ldh dependent glycolytic program in progenitor cells as a means to support growth of the hematopoietic tissue.

Downstream of TCA, event leading to OXPHOS and their loss, as seen in condition with down-regulation of *SdhA* in *Gat^RNAi^* expressing animals (*SdhA^RNAi^; Gat^RNAi^*), which recovered ROS and corrected lymph gland growth (Fig. 4D, E and I-K), did not show recovery in the levels of pPDH and pPDK. Their levels remained comparable to that detected in *Gat^RNAi^* only background (Fig. 6K, L and Supp. Fig. 4E, F). These data with *SdhA^RNAi^; Gat^RNAi^* genetic combination, unlike *Hph^RNAi^; Gat^RNAi^* condition, showed SDH function in moderating ROS generation at the level of OXPHOS and this did not regulate PDH activity status. Thus overall, the data confirmed GABA dependent Hph regulation to maintain PDK activity in progenitor cells. This suppressed PDH activity and controlled TCA cycle, leading to moderation in OXPHOS and subsequent ROS homeostasis in progenitor cells. Additionally, GABA dependent Hph regulation, also controlled Sima and Ldh function in progenitor cells. Together, this contributed to overall growth of the lymph gland.

## Discussion

Central theme of the work undertaken in this study was based on the understanding that ROS as a signaling entity is critical for blood stem-progenitor development and maintenance as reported both in invertebrates and vertebrates (Bigarella et al., 2014; Harris et al., 2013; Prieto-Bermejo et al., 2018; Takubo et al., 2013; Tothova et al., 2007; Vincent & Crozatier, 2010). However, to sustain its developmental role, mechanisms controlling ROS levels that are critical towards its functioning in myeloid progenitor cells remain to be characterized.

Here, we investigated GABA catabolism in lymph gland development and identified a prominent function for GABA breakdown in maintaining progenitor TCA activity. We show progenitors cells of the lymph gland rely on TCA and OXPHOS to generate intracellular ROS. Interestingly, to control their ROS production, progenitor cells internalize extracellular GABA and its breakdown controls PDK phosphorylation. PDK, a key kinase, functions to limit TCA activity, by regulating PDH phosphorylation. This inhibits PDH activity, thereby, blocking the conversion of pyruvate to acetyl-CoA leading to the shutdown of TCA and OXPHOS (Gray et al., 2014; Harris et al., 2002; Wang et al., 2016). Our study demonstrates that the breakdown of GABA into succinate inhibits Hph function. This maintains PDK activity in progenitor cells and consequently lower TCA activity and moderation in progenitor ROS generation. GABA breakdown independent of activating PDK, also stabilizes HIFα/Sima function. Sima controls Ldh, which typically is known to activate a glycolytic state by promoting conversion of pyruvate to lactate (Krejcova et al., 2019; Wang et al., 2016). Thus, metabolism of GABA in progenitor cells allows the maintenance of ROS homeostasis and a metabolic state that drives overall growth of the hematopoietic organ and progenitor homeostasis (Fig. 6M).

### TCA activity and ROS levels in progenitor cells

The MZ progenitor cells, like the mammalian common myeloid progenitor cells maintain elevated ROS. The developmental requirement for ROS is evident in progenitor maintenance and differentiation. Our results show the importance of TCA activity and OXPHOS in the generation of ROS, detected in progenitor cells during homeostasis. Expression analysis of PDH and PDK activity status highlight their regulation and reveal the importance of pyruvate oxidation driving TCA activity in progenitor cells. Loss of PDK function data proves PDK importance in progenitor cell ROS homeostasis and overall growth of the blood tissue.

However, loss of either *Pdha* or *SdhA* function in progenitor cells, that would result in low TCA and low OXPHOS, failed to show any dramatic changes in progenitor homeostasis or overall lymph gland growth. Even though, these genetic conditions led to a reduction in progenitor ROS, these did not show any striking changes in blood progenitor development based on the parameters used in the study. This was unexpected as previously published literature have implicated requirement for ROS in sensitizing progenitor cells to differentiation signals (Owusu-Ansah & Banerjee, 2009). Reduction of ROS, undertaken using over-expression of ROS scavenging enzymes in progenitor cells, impacted their ability to differentiate (Louradour et al., 2017; Owusu-Ansah & Banerjee, 2009). This was however not the case with lowering progenitor TCA activity undertaken here. We predict the independence in differentiation status seen in this study could be an outcome of different genetic approaches utilized to down-regulate ROS (scavenging as opposed to TCA modulation undertaken here). The data also imply fundamental role for additional sources of ROS governing progenitor development and specificity of these pathway in controlling diverse aspects of blood development. This is a reasonable possibility, given the partial reduction in ROS detected upon loss of PDH or SDH function. Duox and Nox dependent mechanisms are other possible sources of cellular ROS (Rada & Leto, 2008) and their effect on blood development should therefore be tested.

### Modulation of PDK activity via Hph but independent of Hifα/Sima

The unexpected finding from the study is the regulation of PDH and PDK function by Hph, which is independent of Sima. In general, the inhibition of PDH activity is considered to be a downstream event of HIFα stabilization (Kim et al., 2006; Papandreou et al., 2006), which is mediated by the transcriptional induction of PDK expression. However, in our study loss of Sima from progenitor cells, a functional ortholog of mammalian Hifα, did not affect expression or activity of either PDH or PDK. On the contrary, our study identifies Hph function as a regulator of PDK activity in blood progenitor cells. Hifα/Sima independent functions of prolyl hydroxylases (Hph) are not unusual, but a mechanistic link between Hph and PDK independent of Hifα /Sima has not been described. The known Hifα independent function of prolyl hydroxylase, include regulation of transcriptional activity of NFκB (Cummins et al., 2006) or Map organizer 1 (Morg1), a WD-repeat protein. Of these, NFκB activation via prolyl hydroxylase regulation is conducted by regulating activity of IKKβ, a kinase that in its non-hydroxylated form phosphorylates an inhibitory factor promoting NFκB activation. A similar mechanism for regulation of PDK activity by Hph can be predicted, but a thorough understanding of this regulation requires further investigation.

### GABA in myeloid metabolism

In addition to ROS, role of GABA in mammalian myeloid cells has gained recent importance because of its ability to metabolically programme these cells (Shao et al., 2021; Steidl et al., 2004; Zhu et al., 2019). In trained innate immune cells, GABA mediated stabilization of Hifα and glycolytic switch has been described (Tannahill et al., 2013). PDK has also been identified as a metabolic checkpoint in myeloid cell polarization (Min et al., 2019). While these studies indicate metabolic commonalities between myeloid system of *Drosophila* and mammals, a comprehensive understanding of metabolic programs via these intermediates during development remained elusive. The work presented in this study highlights dual modes of GABA metabolic function in progenitor cells of the *Drosophila* larvae. GABA via, inhibiting Hph, controls PDK activity and Hifα/Sima stabilization. This mechanism controls ROS balance and glycolytic program in progenitor cells.

PDK activity in progenitor cells can be predicted as a central switch necessary for hematopoietic growth and differentiation. PDK as a regulator of tumor growth is evident in hypoxic conditions (DeBerardinis et al., 2008; Takubo et al., 2013). PDK activity shunts pyruvate away from the citric acid cycle into lactate which keeps the hypoxic cell alive and growing (Kim et al., 2006). Thus, regulation of PDK activity in blood-progenitor cells can be envisaged as a core component whose regulation allows pyruvate availability for other processes contributing to growth. By directing pyruvate utilization into lactate via Ldh and other processes like the pentose phosphate pathway, lymph gland growth can be supported. By invoking on PDK axis, GABA limits pyruvate’s availability for its oxidation via the TCA. This sustains ROS homeostasis more efficiently and allows pyruvate availability for other metabolic arms necessary for blood development and growth.

The current study highlights use of GABA by blood cells to moderate Hph activity and impart it with dual capacity. One being Sima dependent activation of Ldh function and the other being Sima independent, activation of PDK function to limit TCA and OXPHOS in progenitor cells. The common metabolic dependencies seen in mammalian CMPs and *Drosophila* progenitor cells, lends us to hypothesize a similar role for GABA in CMPs which warrants further investigation. The importance of GABA as a central regulator of myeloid development and function across systems including higher mammals is likely to emerge from such studies.

## Material and Methods

### *Drosophila* husbandry, stocks, and genetics

The following *Drosophila* stocks were used in this study: *w^1118^* (wild type*, wt*), *domeMESO-Gal4, UAS-GFP* and *TepIV-Gal4, UAS-mCherry* (Banerjee lab), *Hml^△^-Gal4, UAS-2xEGFP* (S. Sinenko), *UAS-Hph* (C. Frei). The *RNAi* stocks were obtained either from VDRC (Vienna) or BDSC (Bloomington) *Drosophila* stock centres. The lines used for this study are: *Gat^RNAi^* (BDSC 29422)*, Ssadh^RNAi^* (VDRC 106637, BDSC 55683)*, SdhA^RNAi^* (VDRC 330053), *sima^RNAi^* (BDSC 33894)*, Hph^RNAi^* (VDRC 103382)*, Pdha^RNAi^* (BDSC 55345), *Pdk^RNAi^* (BDSC 28635, 35142), *Ldh^RNAi^* (BDSC 33640), *Catalase* (BDSC 24621) and *Sod2^RNAi^* (BDSC 24489). All fly stocks were reared on corn meal agar food medium with yeast supplementation at 25°C incubator unless specified. Tight collections were done for 4-6 hours to avoid over-crowding and for synchronous development of larvae. The crosses involving RNA*i* lines were maintained at 29°C to maximize the efficacy of the *Gal4/UAS RNAi* system. Controls correspond to Gal4 drivers crossed with *w^1118^*.

### ROS (DHE) detection in lymph glands

Lymph glands dissected from the wandering 3^rd^ instar larvae were stained for ROS levels following the protocol of (Owusu-Ansah et al., 2008). The dissected lymph gland tissues were stained in 1:1000 DHE (Invitrogen, Molecular Probes, D11347) dissolved in 1X PBS for 15 min. Tissue were washed in 1X PBS twice and fixed with 4% formaldehyde for 6-8 min. at room temperature in the dark. Tissues were again quickly washed in 1X PBS twice and then mounted in Vectashield (Vector Laboratories). The lymph glands were imaged immediately. A minimum of five independent biological replicates were analysed from which one representative image (one lymph gland lobe) is shown.

### Immunostaining and immunohistochemistry

Immunohistochemistry on lymph gland tissues were performed with the following primary antibodies: mouse-αP1 (I. Ando, 1:30), mouse-αPDH (ab110334, 1:250), mouse-αPDK (ab110025, 1:500), rabbit-αpPDK (TYR243, SAB #11597, 1:200) and rabbit-αpPDH (S293, ab177461, 1:250). The secondary antibodies Alexa Flour 546 and 647 (Invitrogen) were used at 1:500 dilutions. Nuclei were visualized using DAPI (Sigma). Samples were mounted with Vectashield (Vector Laboratories).

Lymph glands dissected from wandering 3^rd^ instar larvae were stained following the protocol of Jung et al., 2005. Lymph gland tissues from synchronized larvae of required developmental stage were dissected in cold PBS (1X Phosphate Buffer Saline, pH-7.2) and fixed in 4% Paraformaldehyde (PFA) for 40 min. at room temperature. Tissues were then washed thrice (15 min. each wash) in 0.3% PBT (0.3% triton-X in 1X PBS) for permeabilization and were further blocked in 5% NGS, for 45 min at RT. Tissues were next incubated in the respective primary antibodies with appropriate dilution in 5% NGS overnight at 4°C. After primary antibody incubation, tissues were washed thrice in 0.3% PBT for 15 min each. This was followed by incubation of tissues in respective secondary antibodies for 2-3 hrs at RT. After secondary antibody incubation, tissues were washed in 0.3% PBT for 15 min. following a DAPI+0.3% PBT wash for 15 min. Excess DAPI was washed off by a wash of 0.3% PBT for 15 min. Tissues were mounted in Vectashield (Vector Laboratories) and then imaged utilizing confocal microscopy. A minimum of ten independent biological replicates were analysed from which one representative image (one lymph gland lobe) is shown.

### Imaging and image processing

DHE stained (ROS) and immuno-stained lymph gland tissues images were acquired using Olympus FV3000 Confocal Microscopy 40X oil-immersion objective. Microscope settings were kept constant for each sample in every experiment. Lymph gland images were processed using ImageJ (NIH) and Adobe Photoshop CS5 software.

### Quantification of lymph gland phenotypes

All images were quantified using ImageJ (NIH) software. Images were acquired as z-stacks and quantifications were done as described previously (Shim et al., 2012). For lymph gland area analysis, middle two z-stacks were merged, and total lymph gland area was marked using the free-hand tool of ImageJ and then analysed further for quantifications. The relative fold change in ROS levels and the intensities per lobe was calculated using mean intensity values. ROS quantifications were done from the entire LG lobe and for intensity quantifications only the dome^+^ (blood progenitor cells) area was marked and then quantified following the protocol of(Madhwal et al., 2020). The area covering the entire LG lobe (for ROS) and dome^+^ region (for intensity quantifications) was marked, and mean intensity was calculated. Background noise was quantified from the unmarked zones at four random regions (marked by equal sized square boxes) and subtracted from the mean intensity values. The relative fold change was calculated from the final mean intensity values and plotted. For all intensity quantifications, the laser settings for each individual experimental set-up were kept constant and controls were analysed in parallel to the mutant conditions every time.

For lymph gland differentiation analysis, middle stack from each image was taken and the LG lobe was marked separately for dome^+^, P1^+^, dome^+^P1^+^ and total area by selecting the respective channels in ImageJ software. Percentage populations from these areas was calculated by dividing each population with total lymph gland area. To calculate percentage negative area, dome^+^P1^+^ area was subtracted from total LG area. dome^+^P1^+^ overlap area was calculated by subtracting dome^+^P1^+^ area from the addition of separate dome^+^ and P1^+^ areas.

### Metabolite supplementation

Succinate (Sodium succinate dibasic hexahydrate, Sigma, S9637) and N-Acetyl-L-cysteine (NAC, Sigma, A7250) enriched diets were prepared by supplementing regular fly food with weight/volume measures of succinate and NAC to achieve 3% and 0.1% concentrations, respectively. Eggs were transferred in these supplemented diets and reared until analysis of the respective tissues (lymph gland).

### Statistical analyses

All statistical analyses were performed using GraphPad Prism nine software and Microsoft Excel 2016. The means were analysed with unpaired *t*-test, two-tailed and medians were analysed with Mann-Whitney test.

## Acknowledgements

We thank I. Ando for the P1 antibody and Bloomington Drosophila Stock Center (BDSC), Vienna Drosophila Resource Center (VDRC) and FlyBase for fly stocks. We acknowledge NCBS, CCAMP for CIFF and fly facility. We thank Dasaradhi Palakodeti and inStem colleagues for helpful discussion and comments on the manuscript. Due to space limitations, we apologize to our colleagues whose work is not cited. This study was supported by the DBT-Center of Excellence grant BT/PR13446/COE/34/30/2015, DST-ECR ECR/2015/000390, 000390DBT-IYBA 2017, CEFIPRA and DBT Ramalingaswami Re-entry Fellowship to TM. MG is a Graduate Student at inStem, in the Mukherjee lab and supported by DST-INSPIRE fellowship.

## Supplementary Figure Legends

**Supplementary Fig. 1 GABA catabolism in *Drosophila* blood progenitor cells controls lymph gland growth.**

DNA is marked with DAPI (blue). Asterisks mark statistically significant differences (*p<0.05; **p<0.01; ***p<0.001, ****p<0.0001 and Mann-Whitney test for **A, B and G** is employed). Lymph glands are demarcated with white dotted line. Scale bar: 20µm. ‘n’ represents number of lymph gland lobes analysed.

**(A)** Quantifications of lymph gland size in *TepIV-mCherry-Gal4 /+* (control, n=20), *TepIV-mCherry-Gal4; UAS-Gat^RNAi^* (n=20, p<0.0001) and *TepIV-mCherry-Gal4*; *UAS-Ssadh^RNAi^* (n=20, p<0.0001) show reduction lymph gland growth as compared to control.

**(B)** Quantifications of lymph gland size in progenitor specific knock-down of Ssadh, *dome-MESO-Gal4, UAS-GFP/+* (control, n=20) and *dome-MESO-Gal4, UAS-GFP*; *UAS-Ssadh^RNAi^* **(BL55683)** (n=20, p<0.0001) show reduction lymph gland growth as compared to control.

**(C-F)** Representative images showing lymph gland size in Hml^+^ loss of *Gat, sima* and *Ldh*

**(C)** Control (*Hml ^Δ^>GFP>/+*) **(D)** expressing *Gat^RNAi^* (*Hml ^Δ^>GFP>Gat^RNAi^*) does not lead to reduction in lymph gland size, **(E)** expressing *sima^RNAi^* (*Hml^Δ^>GFP>sima^RNAi^*) show a mild reduction in lymph gland size and **(F)** expressing *Ldh^RNAi^* (*Hml ^Δ^>GFP>Ldh^RNAi^*) does not show any reduction in lymph gland size as compared to **(C)** control. For quantifications, refer to **G**.

**(G)** Quantifications of lymph gland size in *Hml^Δ^>GFP>/+* (control, n=20), *Hml^Δ^>GFP>Gat^RNAi^* (n=20, p=0.5468), *Hml^Δ^>GFP>sima^RNAi^* (n=20, p=0.0057) and *Hml^Δ^>GFP>Ldh^RNAi^* (n=20, p=0.8831).

**Supplementary Fig. 2. ROS regulation by GABA shunt pathway in *Drosophila* blood progenitors is important for lymph gland growth.**

DNA is marked with DAPI (blue). MZ corresponds to medullary zone. RF is Regular Food and NAC is N-acetylcysteine. Values are mean ± SD, asterisks mark statistically significant differences (*p<0.05; **p<0.01; ***p<0.001, ****p<0.0001, Student’s t-test for G and Mann-Whitney test for **H, M** is employed). Lymph glands are demarcated with white dotted line and are magnified for clarity of staining. Scale bar: 20µm. ‘n’ represents number of lymph gland lobes analysed.

**(A-F)** Representative lymph gland images showing ROS levels in *Sod2^RNAi^* and *Gat^RNAi^*, *Ssadh^RNAi^* on N-acetylcysteine supplementation (NAC), **(A)** Control (*dome-MESO>GFP/+*) showing higher ROS levels in the blood progenitor cells, **(B)** expressing *Sod2^RNAi^* (*dome-MESO-Gal4, UAS-GFP; UAS-Sod2^RNAi^*) in blood progenitor cells leads to increase in ROS levels. **(D)** NAC supplementation to *dome-MESO-Gal4, UAS-GFP; UAS-Gat^RNAi^* and **(F)** NAC supplementation to *dome-MESO-Gal4, UAS-GFP; UAS-Ssadh^RNAi^* rescues the increased ROS phenotype as compared to **(C)** *dome-MESO-Gal4, UAS-GFP; UAS-Gat^RNAi^* on RF and **(D)** to *dome-MESO-Gal4, UAS-GFP; UAS-Ssadh^RNAi^* on RF, respectively. For quantifications, refer to G.

**(G)** Relative fold change in lymph gland ROS levels in *dome-MESO-Gal4, UAS-GFP*/+ (control, n=10), *dome-MESO-Gal4, UAS-GFP; UAS-Sod2^RNAi^* (n=10, p=0.0019), *dome-MESO-Gal4, UAS-GFP*; *UAS-Gat^RNAi^* (RF, n=10, p=0.005), *dome-MESO-Gal4, UAS-GFP; UAS-Gat^RNAi^* (NAC, n=10, p<0.0001), *dome-MESO-Gal4, UAS-GFP*; *UAS-Ssadh^RNAi^* (RF, n=10, p=0.0001) and *dome-MESO-Gal4, UAS-GFP; UAS-Ssadh^RNAi^* (NAC, n=10, p<0.0001).

**(H)** Quantifications of lymph gland area in *dome-MESO-Gal4, UAS-GFP*/+ (control, n=20), *dome-MESO-Gal4, UAS-GFP; UAS-Sod2^RNAi^* (n=20, p=0.0003), *dome-MESO-Gal4, UAS-GFP*; *UAS-Gat^RNAi^* (RF, n=20, p<0.0001), *dome-MESO-Gal4, UAS-GFP; UAS-Gat^RNAi^* (NAC, n=20, p=0.0067), *dome-MESO-Gal4, UAS-GFP*; *UAS-Ssadh^RNAi^* (RF, n=20, p<0.0001) and *dome-MESO-Gal4, UAS-GFP; UAS-Ssadh^RNAi^* (NAC, n=17, p=0.0012).

**(I-L)** Representative images showing lymph gland size, compared to (I) control (*dome-MESO-Gal4, UAS-GFP*/+), (J) *dome-MESO-Gal4, UAS-GFP*; *UAS-Gat^RNAi^* show reduction in lymph gland size. (K) Over-expressing *Catalase* in *Gat^RNAi^* (*dome-MESO-Gal4, UAS-GFP*; *UAS-Catalase*; *UAS-Gat^RNAi^*) leads to rescue of lymph gland size defect as compared to

**(J)** *Gat^RNAi^*. Over-expressing (L) *Catalase* (*dome-MESO-Gal4, UAS-GFP*; *UAS-Catalase*) shows no change in lymph gland size as compared to (I) control. For quantifications, refer to **Fig. 2N**.

**(M)** Relative fold change in lymph gland area in *dome-MESO-Gal4, UAS-GFP*/+ (control, n=20), *dome-MESO-Gal4, UAS-GFP*; *UAS-Gat^RNAi^* (RF, n=20), *dome-MESO-Gal4, UAS-GFP*; *UAS-Hph^RNAi^*; *UAS-Gat^RNAi^* (n=20, p<0.0001), *dome-MESO-Gal4, UAS-GFP*; *UAS-Catalase*; *UAS-Gat^RNAi^* (n=20, p=0.0010), *dome-MESO-Gal4, UAS-GFP*; *UAS-Gat^RNAi^* (SF, n=20, p<0.0001) and *dome-MESO-Gal4, UAS-GFP*; *UAS-Gat^RNAi^* (NAC, n=20, p=0.0043) in comparison to *dome-MESO-Gal4, UAS-GFP*; *UAS-Gat^RNAi^*.

**Supplementary Fig. 3. Medullary zone ROS contributed by TCA regulates lymph gland growth.**

DNA is marked with DAPI (blue). MZ corresponds to medullary zone. Values are mean ± SD and asterisks mark statistically significant differences (*p<0.05; **p<0.01; ***p<0.001, ****p<0.0001 and Student’s t-test for **K, L** and Mann-Whitney test for **M** is employed). Lymph glands are demarcated with white dotted line and are magnified for clarity of staining. White arrows in I-I” indicates the dome^+^P1^+^ overlapping cells. Scale bar: 20µm. ‘n’ represents number of lymph gland lobes analysed.

**(A-C)** Representative lymph gland images showing PDH (red) expression, **(A)** control (*dome-MESO>GFP/+*) lymph gland showing PDH expression, **(B)** expressing *Pdha^RNAi^* in progenitor cells (*dome-MESO-Gal4, UAS-GFP; UAS-Pdha^RNAi^*) show reduction in medullary zone PDH levels and **(C)** expressing *Pdk^RNAi^* in progenitor cells (*dome-MESO-Gal4, UAS-GFP; UAS-Pdk^RNAi^*) does not show reduction in medullary zone PDH levels as compared to **(A)** control. For quantifications, refer to K.

**(D-F)** Representative lymph gland images showing pPDH (red) expression, **(A)** control (*dome-MESO>GFP/+*) lymph gland showing pPDH expression, **(B)** expressing *Pdha^RNAi^* in progenitor cells (*dome-MESO-Gal4, UAS-GFP; UAS-Pdha^RNAi^*) show reduction in medullary zone pPDH levels and **(C)** expressing *Pdk^RNAi^* in progenitor cells (*dome-MESO-Gal4, UAS-GFP; UAS-Pdk^RNAi^*) also show reduction in medullary zone pPDH levels as compared to **(D)** control. For quantifications, refer to L.

**(G-J’’)** Representative images showing lymph gland growth and differentiation status, **(G-G’’)** control (*dome-MESO-Gal4, UAS-GFP*/+) lymph gland showing **(G, G’)** dome^+^ (green), **(G)** dome^-^P1^-^(blue, DAPI) and **(G, G’’)** P1^+^ (red) population, **(H-H’’)** expressing *Pdha^RNAi^* in progenitor cells (*dome-MESO-Gal4, UAS-GFP; UAS-Pdha^RNAi^*) did not show any change in the overall lymph gland differentiation status except a mild increase in the P1 marker population, **(I-I’’)** expressing *Pdk^RNAi^* in progenitor cells (*dome-MESO-Gal4, UAS-GFP; UAS-Pdk^RNAi^*) leads to small lymph gland size and appearance of **(I)** dome^+^p1^+^ overlap population (marked by white arrows and shown in the inset, I) along with an increase in (I”) P1 population, and **(J-J’’)** expressing *SdhA^RNAi^* in the progenitor cells (*dome-MESO-Gal4, UAS-GFP; UAS-SdhA^RNAi^*) did not show any change in the overall lymph gland differentiation status except a mild increase in the P1 marker population and a mild decrease in dome^+^ population as compared to **(G-G’’)** control. For quantifications refer to **Fig. 5K**.

**(K)** Relative fold change in lymph gland MZ PDH levels in *dome-MESO-Gal4, UAS-GFP*/+ (control, n=15), *dome-MESO-Gal4, UAS-GFP*; *UAS-Pdha^RNAi^* (n=20, p<0.0001), and *dome-MESO-Gal4, UAS-GFP*; *UAS-Pdk^RNAi^* (n=10, p=0.0369).

**(L)** Relative fold change in lymph gland MZ pPDH levels in *dome-MESO-Gal4, UAS-GFP*/+ (control, n=20), *dome-MESO-Gal4, UAS-GFP*; *UAS-Pdha^RNAi^* (n=20, p<0.0001), and *dome-MESO-Gal4, UAS-GFP*; *UAS-Pdk^RNAi^* (n=20, p<0.0001).

**(M)** Quantifications of lymph gland area in *dome-MESO-Gal4, UAS-GFP*/+ (control, n=20), and *dome-MESO-Gal4, UAS-GFP; UAS-Pdk^RNAi^* (BL35142, n=20, p<0.0001) also show reduction in lymph gland growth.

**Supplementary Fig. 4. GABA catabolism controls PDH activity via Hph to control lymph gland growth and regulate ROS homeostasis.**

MZ corresponds to medullary zone. Values are mean ± SD and asterisks mark statistically significant differences (*p<0.05; **p<0.01; ***p<0.001, ****p<0.0001 and Student’s t-test for A-F is employed). ‘n’ represents number of lymph gland lobes analysed.

**(A)** Relative fold change in lymph gland MZ PDH levels in *dome-MESO-Gal4, UAS-GFP*/+ (control, n=20), *dome-MESO-Gal4, UAS-GFP*; *UAS-Gat^RNAi^* (n=20, p=0.0819), and *dome-MESO-Gal4, UAS-GFP*; *UAS-Ssadh^RNAi^* (n=13, p=0.0108).

**(B)** Relative fold change in lymph gland MZ PDK1 levels in *dome-MESO-Gal4, UAS-GFP*/+ (control, n=20), *dome-MESO-Gal4, UAS-GFP*; *UAS-Gat^RNAi^* (n=20, p=0.0621), and *dome-MESO-Gal4, UAS-GFP*; *UAS-Ssadh^RNAi^* (n=12, p=0.0344).

**(C)** Relative fold change in lymph gland MZ pPDH levels in *dome-MESO-Gal4, UAS-GFP*/+ (control, n=20), *dome-MESO-Gal4, UAS-GFP*; *UAS-Gat^RNAi^* (RF, n=20, p<0.0001), *dome-MESO-Gal4, UAS-GFP*; *UAS-Gat^RNAi^* (SF, n=16, p<0.0001), and *dome-MESO-Gal4, UAS-GFP*; *UAS-Ssadh^RNAi^* (RF, n=20, p<0.0001), *dome-MESO-Gal4, UAS-GFP*; *UAS-Ssadh^RNAi^* (SF, n=20, p<0.0001).

**(D)** Relative fold change in lymph gland MZ pPDK levels in *dome-MESO-Gal4, UAS-GFP*/+ (control, n=20), *dome-MESO-Gal4, UAS-GFP*; *UAS-Gat^RNAi^* (RF, n=20, p<0.0001), *dome-MESO-Gal4, UAS-GFP*; *UAS-Gat^RNAi^* (SF, n=20, p=0.0016), and *dome-MESO-Gal4, UAS-GFP*; *UAS-Ssadh^RNAi^* (RF, n=20, p<0.0001), *dome-MESO-Gal4, UAS-GFP*; *UAS-Ssadh^RNAi^* (SF, n=12, p=0.0001).

**(E)** Relative fold change in lymph gland MZ pPDH levels in *dome-MESO-Gal4, UAS-GFP*/+ (control, n=20), *dome-MESO-Gal4, UAS-GFP*; *UAS-Gat^RNAi^* (n=20, p<0.0001), *dome-MESO-Gal4, UAS-GFP; UAS-Hph^RNAi^; UAS-Gat^RNAi^* (n=20, p=0.0002), *dome-MESO-Gal4, UAS-GFP; UAS-SdhA^RNAi^; UAS-Gat^RNAi^* (n=16, p=0.0646), *dome-MESO-Gal4, UAS-GFP*; *UAS-Hph^RNAi^* (n=20, p=0.0068) and *dome-MESO-Gal4, UAS-GFP*; *UAS-sima^RNAi^* (n=20, p=0.0507).

**(F)** Relative fold change in lymph gland MZ pPDK levels in *dome-MESO-Gal4, UAS-GFP*/+ (control, n=20), *dome-MESO-Gal4, UAS-GFP*; *UAS-Gat^RNAi^* (n=12, p=0.0008), *dome-MESO-Gal4, UAS-GFP; UAS-Hph^RNAi^; UAS-Gat^RNAi^* (n=20, p<0.0001), *dome-MESO-Gal4, UAS-GFP; UAS-SdhA^RNAi^; UAS-Gat^RNAi^* (n=13, p=0.7726), *dome-MESO-Gal4, UAS-GFP*; *UAS-Hph^RNAi^* (n=20, p<0.0001) and *dome-MESO-Gal4, UAS-GFP*; *UAS-sima^RNAi^* (n=20, p=0.0950).

## Notes

### Competing Interest Statement

The authors have declared no competing interest.

